# Next generation taxonomy: integrating traditional species description with the holobiont concept and genomic approaches - The in-depth characterization of a novel *Euplotes* species as a case study

**DOI:** 10.1101/666461

**Authors:** Valentina Serra, Leandro Gammuto, Venkatamahesh Nitla, Michele Castelli, Olivia Lanzoni, Davide Sassera, Claudio Bandi, Bhagavatula Venkata Sandeep, Franco Verni, Letizia Modeo, Giulio Petroni

## Abstract

In 1991 Margulis defined holobionts as the assemblage of “two or more organisms, members of different species” which remain associate “throughout a significant portion of the life history”. In recent times, holobionts have been described among many and far-related groups of living beings, such as plants, algae, insects, corals, and even humans. These studies have arisen an increasing interest in different contexts but, to our knowledge, the holobiont concept has not been applied in taxonomy. Here we propose a new approach to modern taxonomy, aimed to integrate the holobiont concept and genomic and bioinformatic analyses with the classical/morphological tools traditionally used in taxonomy. The inclusion of symbiont morphology, and of mitochondrial and symbiont genomes will allow the discipline to move toward what could become the “next generation taxonomy”. As an example of this new paradigm in the characterization of holobionts, we herein provide the taxonomic description of the ciliate protist *Euplotes vanleeuwenhoeki* sp. nov. (Euplotia, Ciliophora) and its bacterial endosymbiont “*Candidatus* Pinguicoccus supinus” gen. nov., sp. nov. (*Opitutae, Verrucomicrobia*). Interestingly, we found that this endosymbiont has an extremely reduced genome (~163 Kbp), which is suggestive of a high integration with the host and represents the first case of such an extreme reduction in *Verrucomicrobia*, and the first case in a protist host.

## Introduction

Starting from the de Bary’s definition of symbiosis (“the living together of two differently named organisms” - de Bary 1879), Lynn Margulis defined the term “holobiont” (Meyer-Abich 1950) as the assemblage of “two or more organisms, members of different species” (bionts), which remain associate “throughout a significant portion of the life history” (Margulis 1991). Thus, a holobiont was defined as a compound of different species that together form a single ecological unit. In recent times, an increasing interest regarding holobionts has arisen, leading to different interpretations of the concept (Rosenberg et al. 2007; Zilber-Rosenberg and Rosenberg 2008) and to different theories on the underlying evolutionary driving forces (Rosenberg et al. 2007, 2009; Zilber-Rosenberg and Rosenberg 2008; Rosenberg and Zilber-Rosenberg 2011; Bordenstein and Theis 2015; Bosch and Miller 2016; Roughgarden et al. 2018). Nevertheless, the subject is still debated, especially regarding the applicability of the “holobiont” definition and the evolutionary implications of the concept and, in particular, on the ‘*boundaries*’ of the holobiont, intended as which associations should be considered within this definition. The visions span from the Mindell definition of highly integrated bionts sharing evolutionary paths, which we could view as ‘holobiont *sensu stricto*’, to the broadest sense, thus including even transient members of the microbiome (Mindell 1992, Casiraghi 2012; Moran and Sloan 2015; Douglas and Werren 2016; Hester et al. 2016; Queller and Strassmann 2016; Skilling 2016).

Holobionts have been described among many and far-related groups of living beings, such as plants (for review see: Vandenkoornhuyse et al. 2015; Sanchez-Canizares et al. 2017), algae (Egan et al. 2012), insects (for review see: Guerrero et al. 2013; Minard et al. 2013; Berlanga and Guerrero 2016), corals (Vezzulli et al. 2013; for review see: Thompson et al. 2015; van de Water et al. 2018), and even humans (Postler and Ghosh 2017). In this landscape, protists are under-represented, even though they are highly diverse and environmentally widespread (Šlapeta et al. 2005; Foissner 2006; Weisse 2008), and known for their association with a wide range of other microorganisms (Molmeret et al. 2005; Horn 2008; Ohkuma 2008; Schmitz-Esser et al. 2010; for review see: Gast et al. 2009; Görtz 2010; Nowack and Melkonian 2010; Schweikert et al. 2013; Scheid 2014).

In our opinion the holobiont concept could be very useful for the description of living beings, introducing a new parameter for their taxonomic description, i.e. the presence of stably associated organisms. Indeed, the presence of these organisms can influence the host, its physiology, and its adaptability to the environment (e.g. Rosati et al. 1999, Grosser et al., 2018, Bella et al., 2016); moreover, in some cases it has been shown to influence also its morphology (Giambelluca and Rosati 1996), the fundamental parameter of traditional taxonomy. The introduction of this novel point of view could contribute in rejuvenating the current state of taxonomy (i.e. the science of defining and naming groups of biological organisms based on shared characteristics and, more recently, based on evolutionary relationships).

Classical taxonomy was exclusively based on morphological-comparative techniques requiring a very high specialization on specific taxa. For this reason, and due to the development and rise of modern molecular tools, in the last decades this discipline has faced a significant period of crisis (Mallet and Willmott 2003; Agnarsson and Kuntner 2007). Lately, traditional taxonomy has been renewed in the so-called integrative taxonomy, which also includes ultrastructural and phylogenetic-molecular analysis (Mallet and Willmott 2003; Walter and Winterton 2007).

Now, in our opinion, it is time for another step forward: the future of taxonomic sciences needs to consider and take advantage from the holobiont concept, pursuing a further multidisciplinary integration with modern available technologies, such as bioinformatics and genomic analyses.

We believe that the holobiont concept can be a useful and innovative way to describe a living system in all its components, using a multidisciplinary approach that could further enrich the present integrative taxonomy.

Indeed, although at the moment not mandatory for the description of an organism, the characterization of obligatory or occasional symbionts, *sensu* de Bary (1879) as well as associated microbial consortium (=microbiome), can be considered as an additional useful descriptor of the state of an organism, potentially influencing its development, physiology, and morphology, as observed in previous studies (McFall-Ngai, 2002; Gilbert *et al.*, 2010, 2015; Pradeu, 2011).

As mentioned above, nowadays, the term holobiont is ambiguously defined, indeed it can span from including only obligate mutualistic symbionts to somehow associated microbial consortia. In our opinion, it would be appropriate to apply the concept of holobiont each time the association between different organisms leads to the creation of a functional unit in which emerging characteristics and properties are not typical of the different parts taken separately. In this context, any symbiosis, mutualistic, neutral, or parasitic, affecting either morphology, physiology, or fitness of the host, would be framed.

Such a novel approach implies the necessity to describe each organism of the holobiont and, therefore, the need to build networks of complementary skills, able to combine bio-taxonomy tools, classical morphology, ultrastructure, molecular phylogeny, genomics, and bioinformatics. The proposed framework has the potential to represent a conceptual and methodological advance in taxonomy. Therefore, we propose to define this updated approach as “next generation taxonomy”.

To exemplify the power of this new taxonomic approach for the characterization of holobionts, we herein present the taxonomic description of the ciliate protist *Euplotes vanleeuwenhoeki* sp. nov. (Euplotia, Ciliophora) and its bacterial endosymbiont “*Candidatus* (*Ca.*) Pinguicoccus supinus” gen. nov., sp. nov. (*Opitutae, Verrucomicrobia*). Ciliates are known to form stable associations with eukaryotic (Graham and Graham 1980; Finlay et al. 1987; Kodama and Fujishima 2012; Fokin et al. 2014; Lanzoni et al. 2016; He et al. 2019) and prokaryotic (Rosati et al. 1999; van Hoek et al. 2000; Ferrantini et al. 2009; Modeo et al. 2013a, b; Gong et al. 2014; Serra et al. 2016; Szokoli et al. 2016; Castelli et al. 2019; Fokin et al. 2019) organisms, and thus represent an ideal case of study for the proposed “next generation taxonomy”. Moreover, to the best of our knowledge, this is the first study addressing the concept of holobionts of protists in general.

In detail, we used the proposed approach combining the requirements of an integrative taxonomic description (Warren et al., 2017), and some novel analyses, such as the host mitochondrial genome characterization, with the genomic study on the endosymbiont. Interestingly, we found that the endosymbiont “*Ca.* Pinguicoccus supinus” has an extremely small genome (~163 kbp), comparable in size to extremely reduced genomes of insect symbionts, making this bacterium the first of this category found in a unicellular host (McCutcheon and Moran 2012; Bennett et al. 2016). The extremely small genome size is suggestive of a high level of integration with the host, further indicating the appropriateness of the use of a unifying holobiont/hologenomic approach to ensure a suitable functional and taxonomical description of all the partners involved in such kind of symbioses.

## Material and Methods

### Sample collection and cell culturing

The strain KKR18_Esm was collected on August 8^th^ 2014, in one emissary of the freshwater Lake of Kolleru (16°36′ 05.0″N, E081 18 47.8). Kolleru Lake is the largest freshwater body of India, and a protected Ramsar area due to seasonal migratory birds. It is situated between the deltas of two major rivers, Godavari and Krishna, 15 km south-east from the city of Eluru, in Andhra Pradesh state. The depth of Kolleru Lake is usually around 150-300 cm, but at the time of sampling, the lake was almost dried up and the water level had dropped to the depth of 60-90 cm. Samples were collected at a depth of 15-30 cm near the shore of the lake using sterile 50 ml Falcon tubes: both water and sediment were collected at once.

The original sample was screened by pouring about 20 ml of water in a Petri dish. Single cells were collected using a micropipette, washed several times in mineral water, put in a depression slide and enriched with a few drops of the monoclonal culture of *Dunaliella tertiolecta* (original salinity 5‰, diluted to freshwater) as food to obtain monoclonal cultures. These were maintained in incubator at a temperature of 19 ± 1 °C and on a 12:12 h irradiance of 300 µmo photons/m^2^/s, and progressively adapted to 2.5‰ salinity by means of regular feeding (i.e. once a week) on *D. tertiolecta*.

### Live observations

Live ciliates were observed for morphological identification using differential interference contrast (DIC) microscope with a Leitz Orthoplan microscope (Weitzlar, Germany), with the help of a compression device (Skovorodkin 1990) in order not to distort them as much as possible. For examination of the swimming behaviour, ciliates were observed in a Petri dish, under a stereomicroscope (WILD HEERBRUGG, Switzerland).

### Silver and Feulgen stainings

Ciliates were treated for silver staining analysis with Champy’s solution and then with silver nitrate according to Corliss (1953), to stain the ciliary pattern. Feulgen staining procedure was performed to reveal the nuclear apparatus, using a protocol modified from Dragesco and Dragesco-Kernèis (1986), after cell immobilization with celloidin-diethyl ether-alcohol solution.

### Scanning Electron Microscopy (SEM)

The specimens were fixed in 2% OsO_4_ for 40 min and glued on small coverslips (snipped from 1×1 cm size to the size of stub) previously coated with Poly-L-Lysine and subjected to consecutive dehydration in an ethanol series. Samples were critical point dried according to Nitla et al. (2019). Later, these coverslips were fixed onto SEM stubs with carbon conductive tape. Finally, the samples were sputter-coated with gold (Edwards sputter coater S 150B) and analyzed with JSM-5410 scanning electron microscope.

### Transmission Electron Microscopy (TEM)

*Euplotes* cells were fixed in 2.5% glutaraldehyde in 0.1 M cacodylate buffer for 45 min, rinsed in 0.1 M cacodylate buffer and post-fixed in 1.5% aqueous osmium tetroxide in distilled water for 45 min at room temperature. Then cells were dehydrated and embedded in an Epon-araldite mixture as elsewhere described (Modeo et al. 2013a). The blocks were sectioned with an RMC PowerTome X ultra-microtome. Sections were placed on copper grids and stained with uranyl acetate and lead citrate. Samples were visualized using a JEOL JEM-100SX electron microscope.

### Measurements and recordings

Morphometric data of properly oriented cells were taken by using both live and stained specimen preparations (i.e. Feulgen, silver staining, SEM).

Optical microscopy pictures were captured with a digital camera (Canon Power Shot S45) and used to obtain dimensions of living and stained ciliates. Morphometric measurements were analyzed with ImageJ 1.46r software (Ferreira and Rasband 2012).

Based on micrographs of living and stained cells, accurate schematic line drawings were produced with a procedure described by Montesanto (2015), which employed bitmap graphics with the GNU Image Manipulation Program (GIMP).

Terminology and systematics are mainly according to Curds (1975), Berger (2006), and Lynn (2008).

### DNA extraction and 18S rRNA gene sequencing

Approximately 100-150 cells of KKR18_Esm strain were individually washed 3-5 times in sterile distilled water and fixed in 70% ethanol. Total genomic DNA extraction was performed using the NucleoSpin™ Plant II DNA extraction kit (Macherey-Nagel GmbH and Co., Düren NRW, Germany).

Polymerase chain reaction (PCR) was performed in a C1000™ Thermal Cycler (Bio-Rad, Hercules, CA). The almost full-length of the 18S rRNA gene of *Euplotes* was amplified using the primer combination listed in Supplementary Table 1. High-fidelity Takara Ex Taq PCR reagents were employed (Takara Bio Inc., Otsu, Japan) according to the manufacturer’s instructions. PCR cycles were set as follows: 3 min 94 °C, 35×[30 s 94 °C, 30 s 55 °C, 2 min 72 °C], 6 min 72 °C. PCR products were purified with the Eurogold Cycle-Pure Kit (EuroClone, Milan, Italy) and subsequently sent for direct sequencing to an external sequencing company (GATC Biotech AG, European Custom Sequencing Centre, Germany) by adding appropriate internal primers (see Supplementary Table 1).

### Whole genome amplification and assembly

Starting from around 5-10 cells, the total DNA material was amplified via whole-genome amplification (WGA) method, using REPLI-g Single Cell Kit (QIAGEN^®^, Hilden, Germany). The cells of KKR18_Esm strain were washed in distilled water for three times and the last time in PBS buffer. Then, they were transferred in a 0.2 ml eppendorf together with 4 µl of PBS. The WGA protocol was completed following the manufacturer’s instructions. The so obtained DNA material was processed with a Nextera XT library and sequenced at Admera Health (South Plainfield, USA), using Illumina HiSeq X technology to generate 75,510,798 reads (paired-ends 2×150 bp). Preliminary assembly of resulting reads was performed using SPAdes software (v 3.6.0) (Bankevich et al. 2012).

### Mitochondrial genome assembly and annotation

Contigs representing mitochondrial genome were identified using the Blobology pipeline (Kumar et al. 2013), and by tblastn searches using as queries proteins from reference genomes downloaded from NCBI, namely *Oxytricha trifallax* (JN383842) and *Euplotes minuta* (GQ903130). Contigs with a GC content comprised between 19% and 30%, and a coverage higher than 1000X were selected and a subset of the extracted reads were assembled with SPAdes. We decided to use approximately 10% of the extraxted reads to artifically reduce the reads coverage, as coverage above 100x generally produces worse assemblies, due to an increasing number of exactly replicated sequencing errors, which create false branches in the deBruijn graph. The assembled genome was annotated using PROKKA 1.10 (Seemann 2014), setting the DNA translation codon table “4” and then manually checked.

### Endosymbiont genome assembly and annotation

The presence of symbionts and host related microbial consortium was inspected in the preliminary assembly, using Barrnap (Seeman 2013) to detect 16S rRNA gene sequences, and manually checking all the contigs annotated as bacterial. This, in conjunction with the use of the Blobology pipeline as indicated above, allowed to select the contigs with a GC content lower than 25% and a coverage higher than 1000X for the assembly of the symbiont genome, and a subset (about 10% of the extracted reads. See mitochondrial genome assembly and annotation in Materials and Method section) of the extracted reads were re-assembled with SPAdes. Closure of the circular genome was confirmed via PCR. Specific primers were designed on the basis of genome assembly and used for PCR amplification [Pingui_F162297 (5’ - GTT GTA GCT CTC GGA TCG – 3’), Pingui_R436 (5’ - GTA GAG CAT CTT CGA CTC G – 3’)]; following internal primers were used for sequencing PCR products [Pingui_F163091 (5’ - CTC AGA GCA CTC TGA GAT AG – 3’); Pingui_R199 (5’ - GTT TAG CTC TTC CGA GAT CG – 3’)]. PCR cycles were set as follows: 3 min 94 °C, 35×[30 s 94 °C, 30 s 55 °C, 2 min 72 °C], 6 min 72 °C. We relied on the same reagents, instruments and sequencing company cited above.

The assembled genome was annotated using PROKKA 1.10 (Seemann 2014), setting the DNA translation codon table “4” and then manually checking the results. The predicted protein-coding genes were also classified using NCBI COGs (Galperin et al. 2017), and compared with selected small genomes (Supplementary Table 2) and previously characterized *Verrucomicrobia* genomes (Supplementary Table 3). The COGs thus obtained were used to carry out a Principal Component Analysis (PCA), using SciPy packages in Python, taking into account the numerosity of each COGs class.

### Phylogenetic and phylogenomic analyses

The 18S rRNA gene of the host and the 16S rRNA gene of the newly characterized symbiont were aligned with the automatic aligner of the ARB software package version 5.5 (Westram et al. 2011) on the SSU ref NR99 SILVA database (Quast et al. 2013).

For the analysis on the host, 53 18S rRNA sequences of other members of the *Euplotes* genus, plus 7 sequences of other Spirotrichea as outgroup, were selected (dataset 1).

For the analysis on the symbiont, 98 16S rRNA sequences of other members of *Verrucomicrobia* were selected, plus 12 other sequences belonging to the superphylum *Planctomycetes, Verrucomicrobia, Chlamydiae* (PVC) (Wagner and Horn 2006) as outgroup (dataset 2). Sequences not shown in the tree are listed in (Supplementary Table 4).

After manual editing to optimize base pairing in the predicted rRNA stem regions in each dataset, the two alignments were trimmed at both ends to the length of the shortest sequence. A positional filter was applied to dataset 1, to keep only those columns where the most conserved base was present in at least 10% of the sequences. Resulting matrices contained respectively 1,843 (dataset 1) and 1,456 (dataset 2) nucleotide columns, which were used for phylogeny and for the identity matrix calculation.

For each phylogenetic dataset, the optimal substitution model was selected with jModelTest 2.1 (Darriba et al. 2012) according to the Akaike Information Criterion (AIC). Maximum likelihood (ML) trees were calculated with the PHYML software version 2.4 (Guindon and Gascuel 2003) from the ARB package, performing 1,000 pseudo-replicates. Bayesian inference (BI) trees were inferred with MrBayes 3.2 (Ronquist et al. 2012), using three runs each with one cold and three heated Monte Carlo Markov chains, with a burn-in of 25%, iterating for 1,000,000 generations.

For the phylogenomic analysis, 67 *Verrucomicrobia* (the endosymbiont plus 66 complete genomes from the genome taxonomy database (GTDB) (Parks et al. 2018)), and 4 other members of the PVC superphylum as outgroup, were used. A set of pre-aligned 120 single copy markers were employed (Parks et al. 2018), and the “*Ca.* Pinguicoccus supinus” orthologs were identified by blastp search, then added to the existing alignments with MAFFT v7.123b (Katoh et al. 2013) and concatenated. A positional filter was applied to the concatenated genes according to Parks et al (2018), to obtain a total of 34,747 sites. The best substitution model was estimated with ProtTest (Darriba et al. 2011). RaxML (Stamatakis et al. 2014) was used to estimate ML phylogeny with 1000 bootstraps.

### Fluorescence microscopy

*Euplotes* specimens were fixed for Fluorescence *In Situ* Hybridization (FISH) experiments in 2% OsO_4_ and dehydrated after fixation with an increasing ethanol series for 10 min each. Specimens were processed for hybridization experiments according to previous publications (Boscaro et al. 2013a).

Preliminary FISH experiments were carried out using the generic probe for *Verrucomicrobia* EUB338 III (Daims et al. 1999) and the generic probe for *Bacteria* EUB338 (Amann et al. 1990). Only the EUB338 III probe showed slightly positive signal.

After 16S rRNA sequencing, we designed two probes on the 16S rRNA gene of the putative endosymbiont of *Euplotes*, EUB338 VII (5’ - CTG CTG CCA TCC GTA GAT GT – 3’) and Pingui_1174 (5’ - ACT GAC TTG ACG TCA TCC TCA - 3’).

Both probes were tested *in silico* on the Ribosomal Database Project (RDP) (Cole et al. 2009) and SILVA database using TestProbe 3.0 (Quast et al., 2013), allowing 0 mismatches. The sequence of probe EUB338 VII matched 475 bacterial sequences in the RDP database, 19 of them registered as *Verrucomicrobia* and 139 registered as “unclassified bacteria”. While, probe Pingui_1174 matched 220 bacterial sequences, 35 of them registered as *Verrucomicrobia* and 180 registered as “unclassified bacteria”. Sequences of these two new probes were deposited into probeBase database (Greuter et al. 2016).

Those probes were used to perform additional FISH experiments to confirm the presence of that particular species of *Verrucomicrobia* inside the host.

Hybridized slide preparations were observed under a Zeiss AxioPlan fluorescence microscope (Carl Zeiss, Oberkochen, Germany) equipped with an HBO 100W/2 mercuric vapor lamp at the following UV wavelengths: ~ 495 nm, ~ 550 nm. Digital images were captured at different magnifications (40X and 100X) by means of a dedicated software (ACT2U, version 1.0).

### Endosymbiont 16S rRNA gene screening on IMNGS

Diversity and environmental distribution of the bacterial endosymbiont was estimated using the IMNGS on-line platform (Lagkouvardos et al. 2016), which screens most of 16S rRNA gene amplicon datasets available. A query at 95% similarity was performed using the endosymbiont’s full-length 16S rRNA gene and related *Puniceicoccaceae* sequences (accession numbers: AB073978, AB372850, AB614893, AB826705, CP001998, DQ539046, EU462461, KT751307, JQ993599, JQ993517, Y19169, for their 16S rRNA full-length gene phylogenetic position see Figure 7). The obtained sequences were longer than 300 bp and were divided into three different groups to perform phylogenetic analysis according to the 16S rRNA gene hypervariable regions V1-V2, V4-V6, and V7-V8. Sequences were clustered in OTUs with a 99% threshold similarity using UCLUST (Edgar 2010), then they were aligned with MUSCLE (Edgar 2004), and FastTree (Price et al. 2009) was employed to infer phylogenetic analyses. Thereafter, environmental distribution was investigated using the endosymbiont sequence as query and their abundances were calculated.

## Results

### Description of *Euplotes vanleeuwenhoeki* sp. nov. (Figures 1–5, Table 1)

Phylum Ciliophora Doflein, 1901
Class Spirotrichea Bütschli, 1889
Subclass Euplotia Jankowski, 1979
Order Euplotida Small and Lynn, 1985
Family Euplotidae Ehrenberg, 1838
Genus *Euplotes* Ehrenberg, 1831

#### Diagnosis

Size *in vivo* (*X*±SD) 49.1 ± 4.7 × 32.7 ± 3.8 μm. Dorso-ventrally flattened, with an oval to ellipsoidal shape. “C–shaped” or “3-shaped” macronucleus and a single micronucleus. Dargyrome of double-*eurystomus* type, 7-8 dorsal ridges, with 13–14 dikinetids in the mid–dorsal row. About 22-29 adoral membranelles. Cirri pattern: 10 fronto-ventral, 5 transverse, 2 marginal, and 2 caudal cirri. Freshwater.

#### Type locality

Freshwater emissary of Kolleru Lake, in the proximity of Allapadu-Kolletikota road, West Godavari District of Andhra Pradesh, India. This species inhabits freshwater sites covered by *Eichhornia* sp. (water hyacinth).

#### Etymology

We dedicated this new species of *Euplotes* to Antoni Philips van Leeuwenhoek (1632-1723), Dutch optician and naturalist. Van Leeuwenhoek is best known for his pioneering work in microscopy and for his contributions toward the establishment of microbiology as a scientific discipline. For this reason, he is also known as “the father of microbiology”, being one of the first microscopists and microbiologists.

#### Type material

The slide with the silver-stained holotype specimen (indicated with a black circle of ink on the coverslip) and some paratype specimens has been deposited in the collection of the “Museo di Storia Naturale dell’Università di Pisa” (Calci, Pisa, Italy) with registration number “2019-1”. Two slides with silver-stained paratype specimens (indicated with a black circle of ink on the coverslip) were deposited in the collection of the Natural History Museum of London (registration number: NHMUK 2019.3.16.1), and in the collection of the Unit of Zoology-Antropology of the Department of Biology at Pisa University (registration number: UNIPI_2019-1), respectively.

#### Morphological description

Size (*X*±SD) *in vivo* 49.1 ± 4.7 × 32.7 ± 3.8 μm. Size after silver staining 45.6 ± 3.1 × 29.1 ± 4.1 μm. Cell reduction after fixation: 8%. Cells dorso-ventrally flattened, with an oval to ellipsoidal shape (Fig.1). Right margin usually straight or slightly convex, left margin tapered in the anterior, becoming convex in the mid–body, and both ends are rounded (Fig.1a). Ciliates can crawl on the substrate and swim freely in the medium. Cytoplasm transparent with some roundish, yellow granules; few food vacuoles containing green algae and bacteria (Fig. 1b, c). Single contractile vacuole located at the level of transverse cirri (Fig. 1b). On dorsal side, cortical ampules arranged around each bristle form conspicuous rosettes with their cortical insertions (Fig. 1c). Macronucleus (Ma) “C–shaped” or “3-shaped” (size: 36.6 ± 4.5 × 5.7 ± 1.0 μm) with irregularly dense chromatin, and a single, roundish micronucleus (Mi) (diameter: 2.0 ± 0.2 μm), usually located in a small depression close to Ma (Fig. 1d).

**Figure 1.**
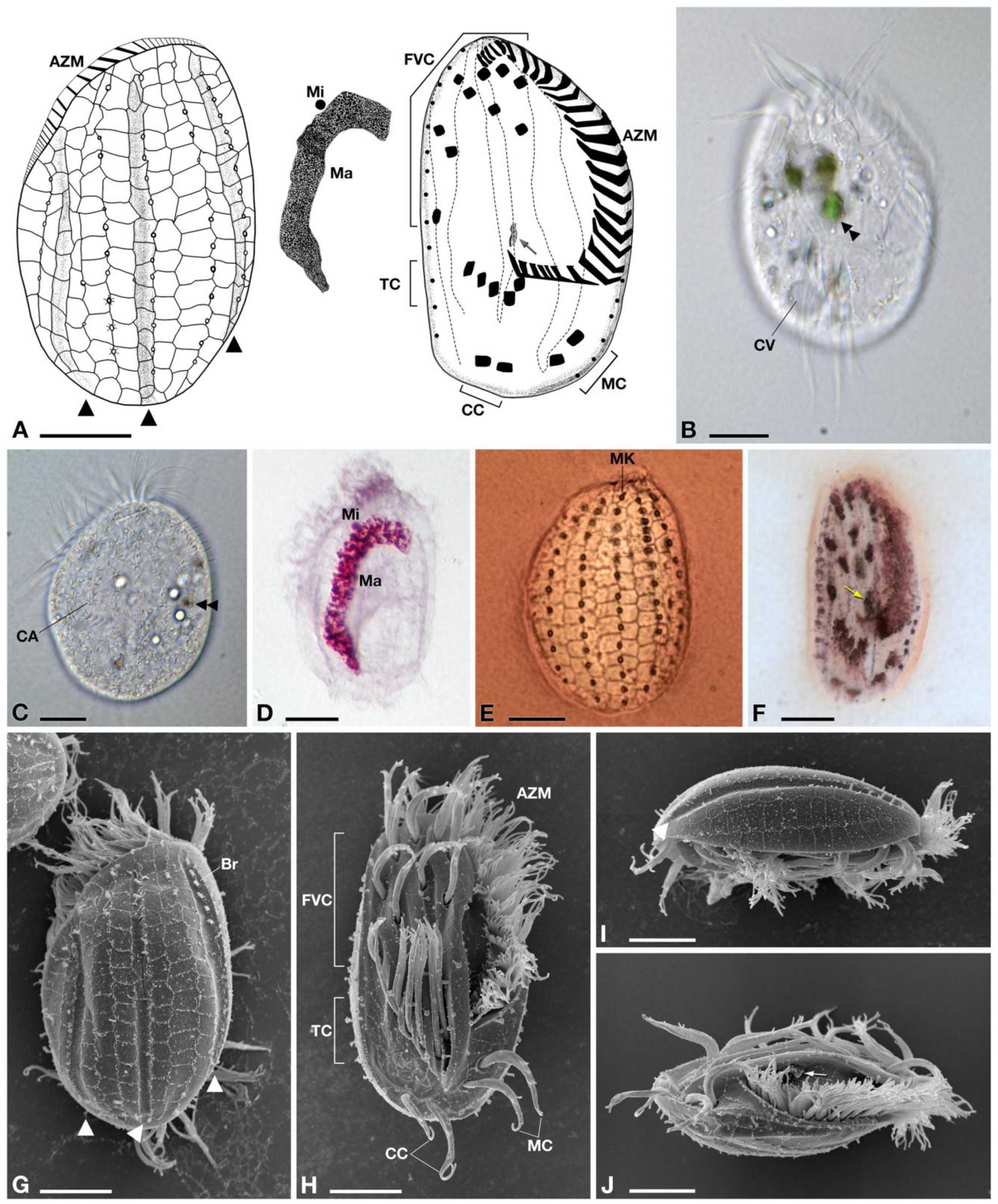
Morphology of *Euplotes vanleeuwenhoeki* sp. nov. A) Schematic drawings of the dorsal side (left), ventral side (right), and nuclear apparatus (middle); B) Live picture, ventral side. C) Live picture, dorsal side; D) Feulgen staining, showing macronucleus (Ma) and micronucleus (Mi); E) Silver staining, dorsal side; F) Silver staining, ventral side; G-J) SEM pictures of dorsal side (G), ventral side (H), and lateral views (I-J); Arrow: paroral membrane; Arrowhead: dorsal furrow; Double arrowhead: food vacuole containing algae; AZM: adoral zone of membranelles; Br: bristle; CA: cortical ampules; CC: caudal cirri; CV: contractile vacuole; FVC: fronto-ventral cirri; Ma: macronucleus; MC: marginal cirri; Mi: micronucleus; MK: mid-dorsal kinety; TC: transverse cirri. *Bars* stand for 10 µm.

Dargyrome of the double–*eurystomus* type, with two rows of polygonal alveoli between each pair of dorsolateral kineties (Figs 1e, g, i). Dorsal surface crossed by three longitudinal furrows (i.e. right marginal, median, and left marginal), reaching the posterior region of the cell (Fig. 1g). Six dorsolateral kineties, three in correspondence of dorsal furrows, carrying short bristle-like cilia (Fig. 1g); the leftmost kinety is placed in a slightly ventrolateral position (Fig. 1j). Mid–dorsal row containing up to 13–14 dikinetids (Fig. 1e).

On the ventral side, invariably 10 frontoventral cirri (FVC), 5 transverse cirri (TC), 2 well developed caudal cirri (CC), and 2 marginal cirri (MC) on the left side, in the posterior end of the cell (Fig. 1f, h, j). Argyrome is highly irregular (Figs 1h, j) and the ventral surface presents five longitudinal ridges hosting cirral insertions; the three ridges on the left are more prominent (Fig. 1h). The first and the fifth ridges reach the posterior part of the cell at level of the CC, while the other three ridges terminate beyond the TC (Fig. 1h).

Peristome narrow, extending for about 63% of the body length, on the ventral side. Adoral zone comprising 22–29 membranelles (AZM), starting at the top of the cell, travelling down along the left side and reaching the first ventral ridge, with a slight curve towards the centre of the body, at level of transverse cirri (Fig. 1h). The paroral membrane appreciable in silver stained specimens (Fig. 1f), and in SEM-processed specimens (Fig. 1j), although carrying cilia shorter than those forming the AZM. All morphometric data are shown in Table 1.

**Table 1.** Morphometric data for *Euplotes vanleeuwenhoeki* sp. nov.

| Character | MIN-MAX | $\bar{X} \pm \text{SD}$ | CV | n |
| --- | --- | --- | --- | --- |
| **Body length ($\mu\text{m}$)** |  |  |  |  |
| Live | 39.5-58.0 | 49.1±4.7 | 9.7 | 22 |
| SI | 38.9-50.3 | 45.6±3.1 | 6.7 | 37 |
| **Body width ($\mu\text{m}$)** |  |  |  |  |
| Live | 24.9-37.8 | 32.7±3.8 | 11.6 | 22 |
| SI | 21.6-35.5 | 29.1±4.1 | 14.0 | 37 |
| **Macronucleus length ($\mu\text{m}$)** |  |  |  |  |
| Feulgen staining | 28.9-48.6 | 36.6±4.5 | 12.4 | 15 |
| **Macronucleus width ($\mu\text{m}$)** |  |  |  |  |
| Feulgen staining | 4.3-7.2 | 5.7±1.0 | 17.3 | 15 |
| **Micronucleus diameter ($\mu\text{m}$)** |  |  |  |  |
| Feulgen staining | 1.8-2.3 | 2.0±0.2 | 9.0 | 15 |

MIN: minimum value; MAX: maximum value; $\bar{X}$ : arithmetic mean; SD: standard deviation; CV: coefficient of variation (%); n: number of specimens analyzed; SI: silver impregnation.

#### Fine structure

The fine structure of *E. vanleeuwenhoeki* (Fig. 2) matches that of the other previously described *Euplotes* species, in general showing typical features (Fauré-Fremiet and Andrè 1968; Nobili and Rosati Raffaelli 1971; Kloetzel 1974; Ruffolo 1976; Modeo et al. 2005; Schwarz et al. 2007). Under the cell cortex flat alveoli are present (Figs 2a-f, h-j). On the dorsal side, somatic cilia consisting of dikinetids (Figs 2a, c) are deeply inserted into the cytoplasm (~ 1.4 µm); from kinetosomes only a single bristle-like cilium emerges (Fig. 1c, g). In the bristle pit, some filamentous material is sometimes visible. (Fig. 2b). This is likely released by cortical ampules, the typical exocytotic organelles associated with both *Euplotes* dorsal bristle and compound ciliary organelles of the ventral surface; these organelles probably represent specialized compartments of the cell in which materials that need to be excreted are accumulated, stored, and released according to the requirements of the different *Euplotes* species (Rosati and Modeo 2003). Ampules associated with dorsal bristles of *E. vanleeuwenhoeki* appear elongated (size: ~ 1.6 x 0.3 µm) and usually empty possibly also due to fixation procedure (Fig. 2c). Membranelles bordering the upper and left side of the oral cavity are separated from each other by ridges (Fig. 2d). Each membranelle of AZM consists of three rows of cilia: two equally long plus a shorter one (Fig. 2d). Axonemes contain many electron dense granules (Fig. 2d). Kinetosomes of membranelles are linked at their base. (Fig. 2e). A polystichomonad paroral membranelle is inserted on the right margin of the terminal oral cavity; its cilia appear linked to each other at the kinetosome level (Figs 2d, e). Many flat, electron lucid pharyngeal disks are associated to the base of the cytostome, in correspondence of AZM bases (Figs 2e, f). Macronucleus contains large piece of chromatin and large nucleoli (Figs 2f, g). Micronucleus consists of fine chromatin (Fig. 2g). A single contractile vacuole with an irregular silhouette is observed near a transverse cirrus (Fig. 2h). On the ventral side, kinetosomes of cilia forming cirri contain large electron dense granules (Fig. 2i). Mitochondria show variable shape and size and typical tubular cristae (Figs 2b, c, e, f). Lipidic reserve substances consist of large granules; polysaccharidic reserve substances are represented by rosettes of glycogen abundantly and sparsely distributed throughout the cytoplasm (Figs 2g, h). Large, irregular phagosomes are also present, with various content in different digestion stages (Fig. 2j).

**Figure 2.**
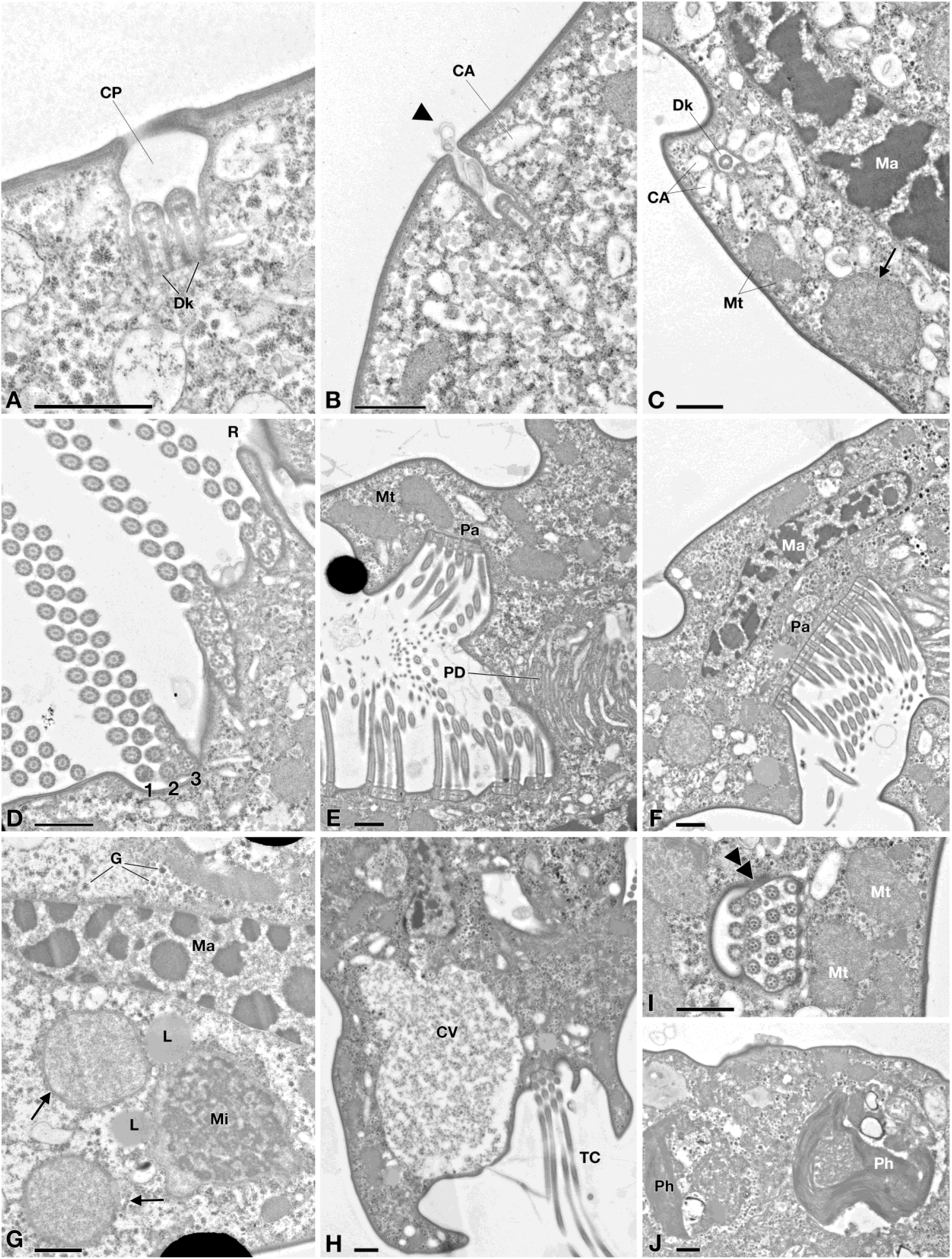
TEM picture of *Euplotes vanleeuwenhoeki* sp. nov. A-C) Cortex region; D-F) Oral region; A) Ciliary pit (CP) with dikinetid (Dk); B) Detail of bristle pit containing filamentous material (arrowhead); cortical ampules (CA) are visible; C) Transverse section of dikinetid (Dk) surrounded by cortical ampules; sections of the macronucleus (Ma), two mitochondria (Mt) and an endosymbiotic bacterium (arrow) are visible; D) Detail of oral membranelles, composed of two longer rows (1, 2) and one shorter row (3) of cilia; membranelles are separated by ridges (R); E) Section of oral region, showing paraoral membrane (Pa) in front of oral membranelles and pharingeal disks (PD); F) Closer view of paraoral membrane and transverse section of macronucleus; G) Endosymbiotic bacterial cells (arrow) inside *Euplotes* cytoplasm, nearby macronucleus and micronucleus (Mi); rosettes of glycogen (G) and lipid droplets (L) are present; L) Detail of contractile vacuole (CV) and transverse cirrus (TC); I) Section of cirrus (double arrowhead), close to mitochondria; J) Two phagosomes (Ph). CA: cortical ampules; CP: ciliary pit; CV: contractile vacuole; Dk: dikinetid; G: rosette of glycogen; L: lipid droplet; Ma macronucleus; Mi micronucleus; Mt: mitochondrion; Pa: paraoral membrane; PD: pharingeal disk; Ph: phagosome; R: ridge; TC: transverse cirrus; Arrowhead: filamentous material; Arrow: endosymbiotic bacterium; Double arrowhead: transverse section of cirrus. *Bars* stand for 1 µm.

Numerous, morphologically similar endosymbiotic bacteria, presenting variable shape and size, are located in the cytoplasm (Figs 2, 5): a detailed morphological description is presented below.

#### Gene sequence

The 18S rRNA gene sequence of *E. vanleeuwenhoeki* (strain KKR18_Esm) obtained from PCR resulted 1,849 bp long, and it has been deposited in NCBI GenBank database with the accession number KY855568. The 18S rRNA gene sequence of *E. vanleeuwenhoeki* showed the highest identity with sequences of *Euplotes* cf. *antarcticus* (FJ998023) and *E. trisulcatus*, (EF690810): 99.0% (3 gaps, 16 mismatches) and 98.7% (13 gaps, 19 mismatches), respectively (Supplementary Table 5).

#### Phylogeny

The 18S rRNA gene-based phylogeny placed *E. vanleeuwenhoeki* in the so-called “clade A” of genus *Euplotes* (Syberg-Olsen et al. 2016; Boscaro et al. 2018), clustering together with *Euplotes* cf. *antarcticus* (FJ998023; Gao and Song unpublished) and with *E. trisulcatus*, (EF690810; Schwarz and Stoeck unpublished), with high statistical support (1.00/100). This clade resulted sister to a clade comprising sequences attributed to *E. charon* (AF492705), *E. magnicirratus* (AJ549210), and *E. euryhalinus* (EF094968, JF903799) group (see later discussion on species attribution). See Figure 3.

**Figure 3.**
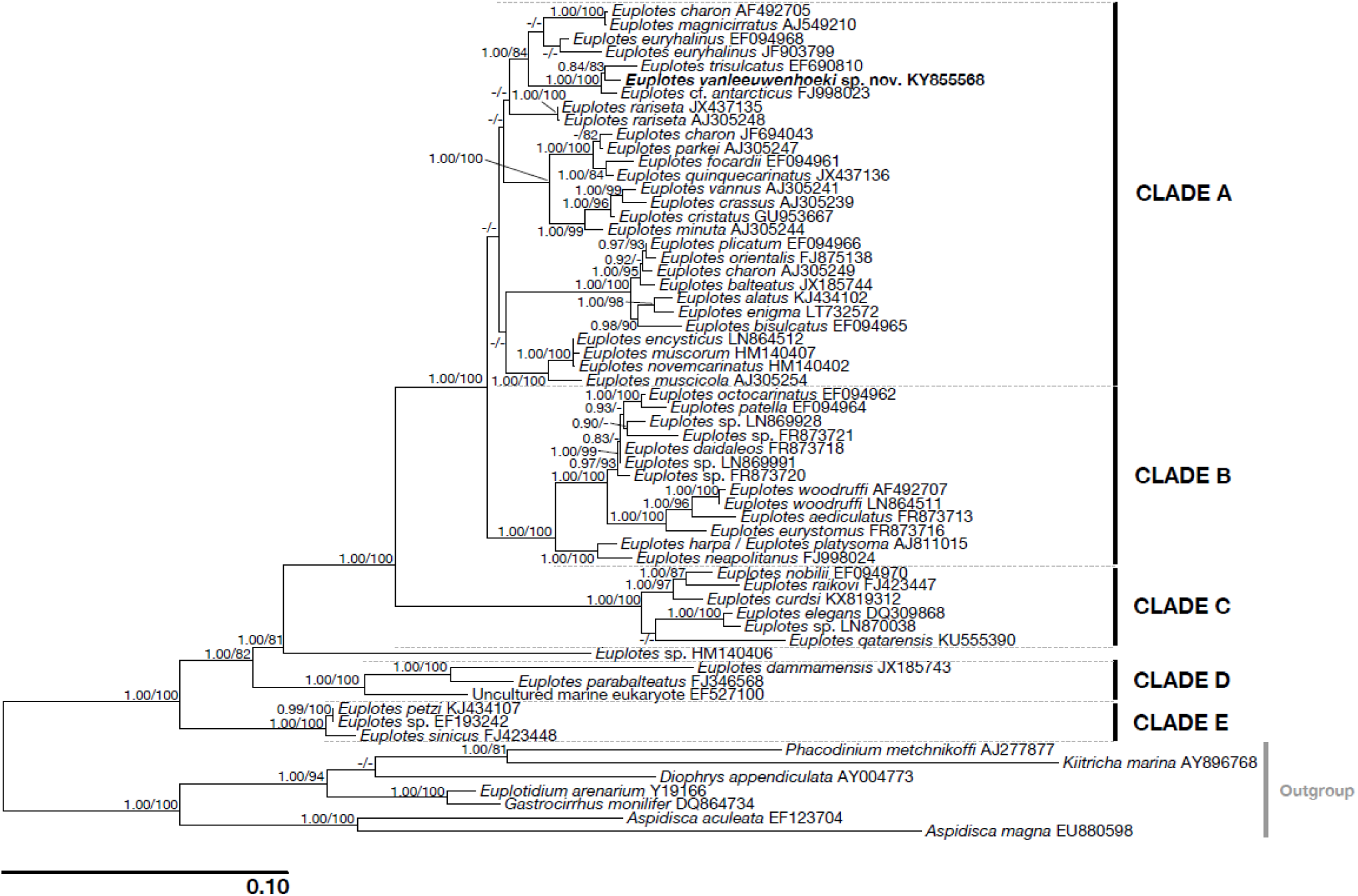
Phylogenetic tree of genus *Euplotes* based on the 18S rRNA gene. Numbers associated to nodes represent posterior probabilities and bootstrap values, respectively (only values above 0.80–75 are shown). Sequence obtained in the present work is in *bold*.

#### Mithocondrial genome

The assembly resulted in a single linear contig 41,682 bp long with a GC content of ~0.25%, representing the complete mitochondrial genome of *E. vanleeuwenhoeki*. It has been deposited in NCBI GenBank database with the accession number MK889230. It contains 36 protein coding genes and 16 tRNAs. The genome presents the 16S rRNA and 23S rRNA genes split in two loci, with the 23S rRNA further divided in two genes, separated by a short interposing region of approximately 350 nucleotides (Fig. 4). The predicted direction of the transcription is away from a central region constituted of low-complexity repeated units (Fig. 4). The splitting of the rRNA genes and the presence of a central repeat region is a common feature shared by all the so far investigated *Euplotes* mitochondrial genomes, (De Graaf et al. 2009) and by the one of *Oxytricha trifallax* (Swart et al. 2011). The novel genome shows an overall synteny with the mitochondrion of of *Euplotes minuta, Euplotes crassus* (De Graaf et al. 2009) and *Oxytricha trifallax* (Swart et al. 2011), with the exception of the two terminal regions, which show a different structure in respect of the other three genomes (Fig. 4).

**Figure 4.**
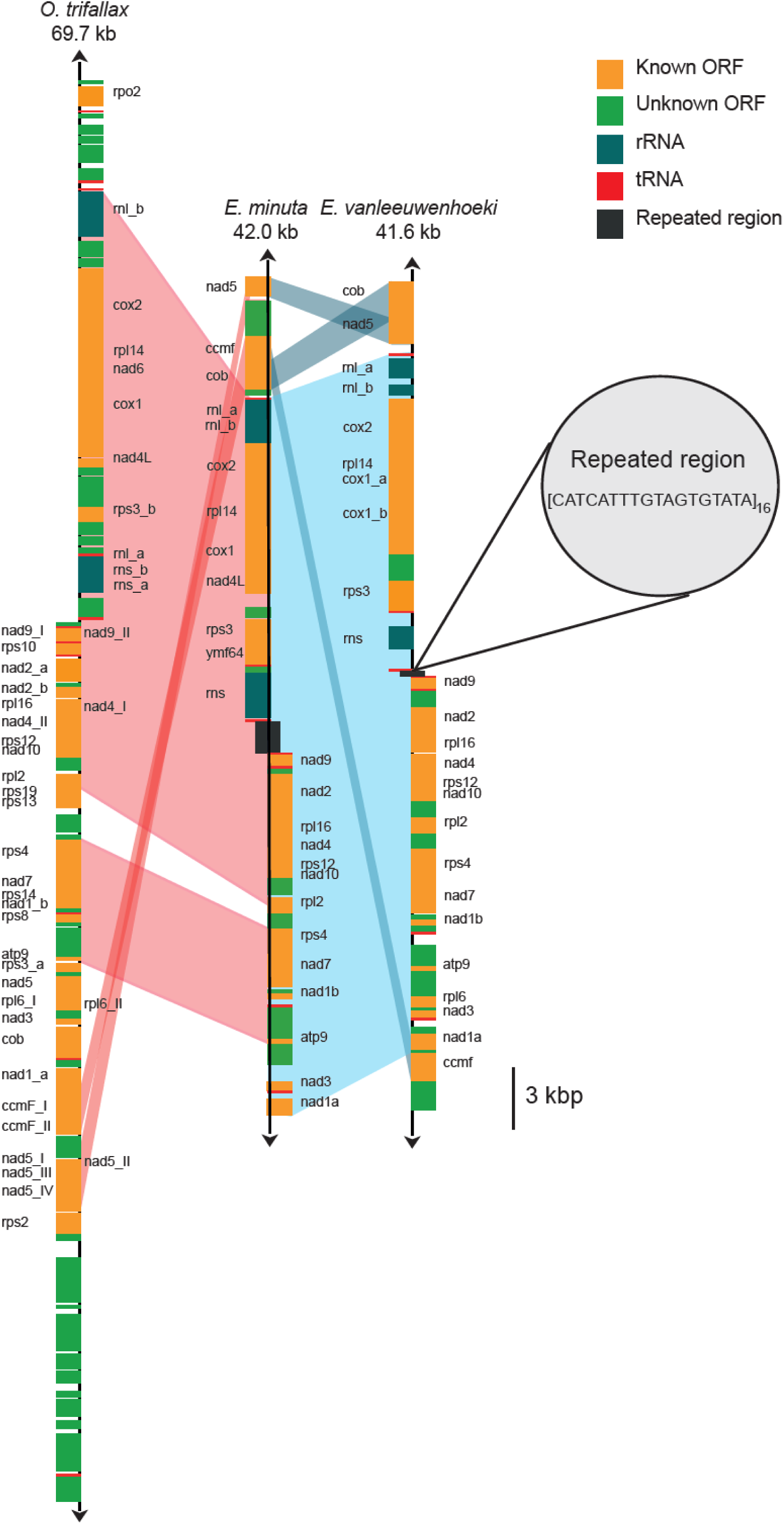
Mitochondrial gene map of *Euplotes vanleeuwenhoeki* sp. nov. The gene map of the mitochondrial genome of *E. vanleeuwenhoeki* in comparison with those belonging to *Euplotes minuta* and *Oxytricha trifallax* is represented. Homologous regions among the three genomes are indicated by pale coloured areas. Names of split genes are suffixed by a letter or a lowercase Roman numeral. The direction of transcription is indicated by an arrow at each end of the mitochondrial map.

#### Microbial consortium

The screening of the preliminary assembly for bacterial 16S rRNA genes allowed to identify the presence of a single microorganism associated to *E. vanleeuwenhoeki*. Further analyses proved that this bacterium was localized in the cytoplasm of the ciliate, and that it was a novel endosymbiont we named “*Ca.* Pinguicoccus supinus” (see “Endosymbiont characterization” section). No other bacterial 16S rRNA gene sequence was detected in the sequencing reads. Moreover, most of the other contigs tha were preliminary flagged as bacterial from the best megablast hit in the Blobology pipeline actually belonged to the mitochondrial or nuclear genome of *Euplotes*, or they were short (< 600 bp) or at a very low coverage (< 30x), thus were considered as from undetermined origin, possibly representing only minor contaminations, and were discarded (Supplementary Table 6).

### Endosymbiont characterization: *“Candidatus* Pinguicoccus supinus*” gen. nov. sp. nov.* (Figures 5–10)

#### Morphological description

“*Ca.* Pinguicoccus supinus” gen. nov. sp. nov. is a roundish-ovoid bacterium detected in the cytoplasm of *E. vanleeuwenhoeki* (Fig. 5) with a diameter of 1.3-2.3 µm (on average (*X*±SD): 1.9 ± 0.3 µm). It usually lies beneath the ciliate cortex, often in clusters of several individuals (Fig. 5a). Although the most common bacterial shape observed is rounded (Fig. 5b), sometimes ovoid (Fig. 5c) and irregular (Figs 5d, e) individuals can be detected as well. This cell shape plasticity might possibly be due to the pressure exerted by the host cytoplasm on the ductile body of the bacterium. No symbiosome is observed to isolate the endosymbiont from ciliate cytoplasm (Fig. 5). “*Ca.* Pinguicoccus supinus” is delimited by a double membrane with a thin space between the two layers possibly corresponding to the paryphoplasm (Fig. 5b), the intracellular space defined for the first time by Linsday and colleagues (Lindsay et al. 2001). In several individuals, the increase of membrane area is visible: in some cases, a slight invagination of the inner membrane occurs (Figs 5b-d), while in others the evagination of the external membrane can be observed (Fig. 5d-f). In the latter case, different inclusions of unknown origin have also been observed in the space between inner and outer membrane (Fig. 5d, e). The bacterial cytoplasm (possibly corresponding to the pirellulosome; Lindsay et al. 2001) generally appears homogeneous and a compact, more electrondense region, likely corresponding to bacterial nucleoid, is visible in some bacteria with an eccentric localization (Fig. 5d, e, h, i). Occasionally, specimens show a very emphasized folding of membrane area, making it difficult to recognize whether the folding comes from the inner or the outer membrane (Fig. 5g). Out of a total of ~ 80 observed endosymbionts observed in thin section, roughly one third are in proximity (i.e. at a distance of ~ 0.25 µm or less) of mitochondria (Figs. 5a-c, f, i) and one fifth are in proximity to lipid droplets (Figs 5e, g, h). Intriguingly, in these cases, bacterial double membrane is even seen somehow in direct contact with mitochondrial external membrane (Fig. 5c) and lipid droplets (Fig. 5h). *“Ca.* Pinguicoccus supinus” reproduces in the host cytoplasm by binary fission and has, apparently, a typical symmetrical division (Fig. 5i).

**Figure 5.**
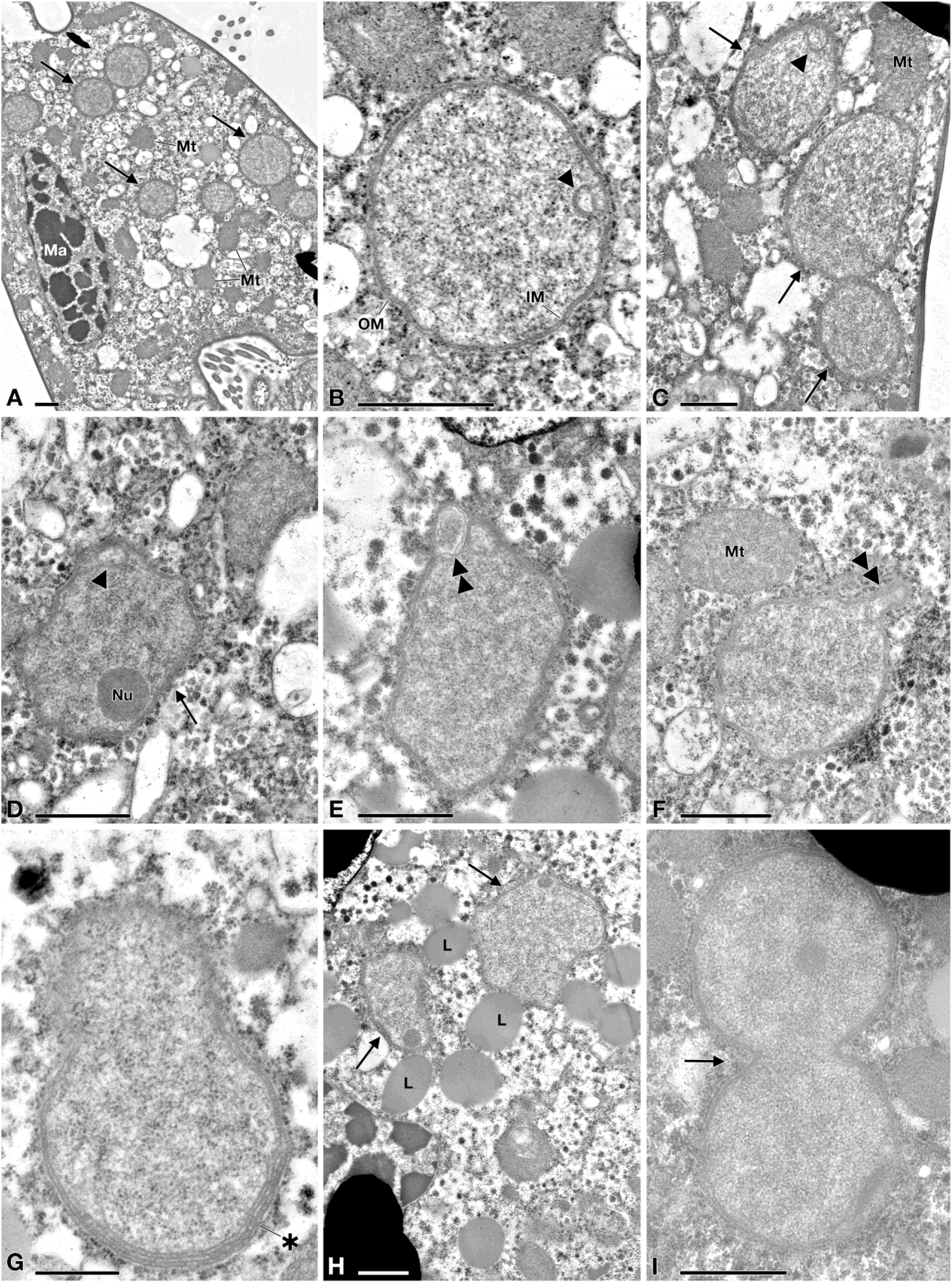
TEM pictures of “*Candidatus* Pinguicoccus supinus”. A) Endosymbiont cells (arrow) in host cytoplasm, lying underneath the cortex, aggregated in clusters; macronucleus (Ma) and some mitochondria (Mt), also in proximity or in apparent close contact with endosymbionts, are visible; B) Closer view of “*Ca.* Pinguicoccus supinus” showing inner membrane (IM) and outer membrane (OM); an invagination of the inner membrane (IM) is present (arrowhead); C) Different cell shapes of three “*Ca.* Pinguicoccus supinus” specimens (arrow), from rounded to ovoid; the inner membrane is invaginated (arrowhead); D) Endosymbiont cell with an irregular shape, with and evident nucleoid (Nu) and an evagination of the outer membrane (double arrowhead); E-F) “*Ca.* Pinguicoccus supinus” cells showing evagination of the outer membrane (double arrowhead); F) Endosymbiont cell in proximity to a mitochondrion; G) “*Ca.* Pinguicoccus supinus” showing emphasized folding (asterisk) of membrane area; H) Endosymbiont cells (arrow) appear to be in close contact with lipid droplets (L); I) “*Ca.* Pinguicoccus supinus” (arrow) during binary fission; the division septum is well visible. IM: inner membrane; L: lipid droplet; Ma: macronucleus; Mt: mitochondrion; Nu: nucleoid; OM: outer membrane; Arrow: “*Ca.* Pinguicoccus supinus” cell; Arrowhead: evagination of the outer membrane; Asterisk: folding of membrane area; Double arrowhead: section of bacterial cell folding. *Bars* stand for 1 µm (A-F, H-I) and 0.5 µm (G).

#### Gene sequence

The 16S rRNA gene sequence of “*Ca.* Pinguicoccus supinus” resulted 1,517 bp long and has been deposited in NCBI GenBank database with the accession number MK569697. It showed highest identity (78.8; 78.3%; 78.2%) with sequences from uncultured bacteria (JQ993517 and MNWT01000005; AB826705; AY571501 respectively). The best hit with a cultured bacterium was 76.8% with *Ruficoccus amylovorans* (KT751307) (*Verrucomicrobia*, *Opitutae*, *Puniceicoccaceae*). In general, identity values with the closest relatives resulted quite low (75.7-78.8%) (Supplementary Table 7).

#### Genome assembly

The assembly of the symbiont’s genome resulted in a single circular chromosome, 163,218 bp long with a GC content of 25.1%. The complete genome sequence of “*Ca.* Pinguicoccus supinus” has been deposited in NCBI GenBank database with the accession number CP039370. It contains 168 protein coding sequences, 34 tRNAs and a single rRNA operon composed by a 16S rRNA gene, a 23S rRNA gene and a 5S rRNA gene. The overall coding percentage is 92.3%. The ORFs were subjected to clusters of orthologous groups (COGs) classification, and 131 COGs were identified, most of which were related to the general cellular function categories (J, O, M and I, for a total of 112 COGs - Supplementary Table 8). This repertoire is a subset of the previously characterized *Verrucomicrobia*, i.e. without any exclusive metabolic pathway or gene (Supplementary Table 3). Considering the drastic genome reduction of “*Ca.* Pinguicoccus supinus”, we decided to perform comparative analyses using as reference the few other symbiotic bacteria with highly reduced genome although belonging to unrelated lineages (i.e. *Bacteroidetes, Alphaproteobacteria, Gammaproteobacteria, Betaproteobacteria*). PCA was able to capture almost 56% of the whole variance (Fig. 6a) in the COG dataset (Component one 32% of explained variance, and Component two 24%).

**Figure 6.**
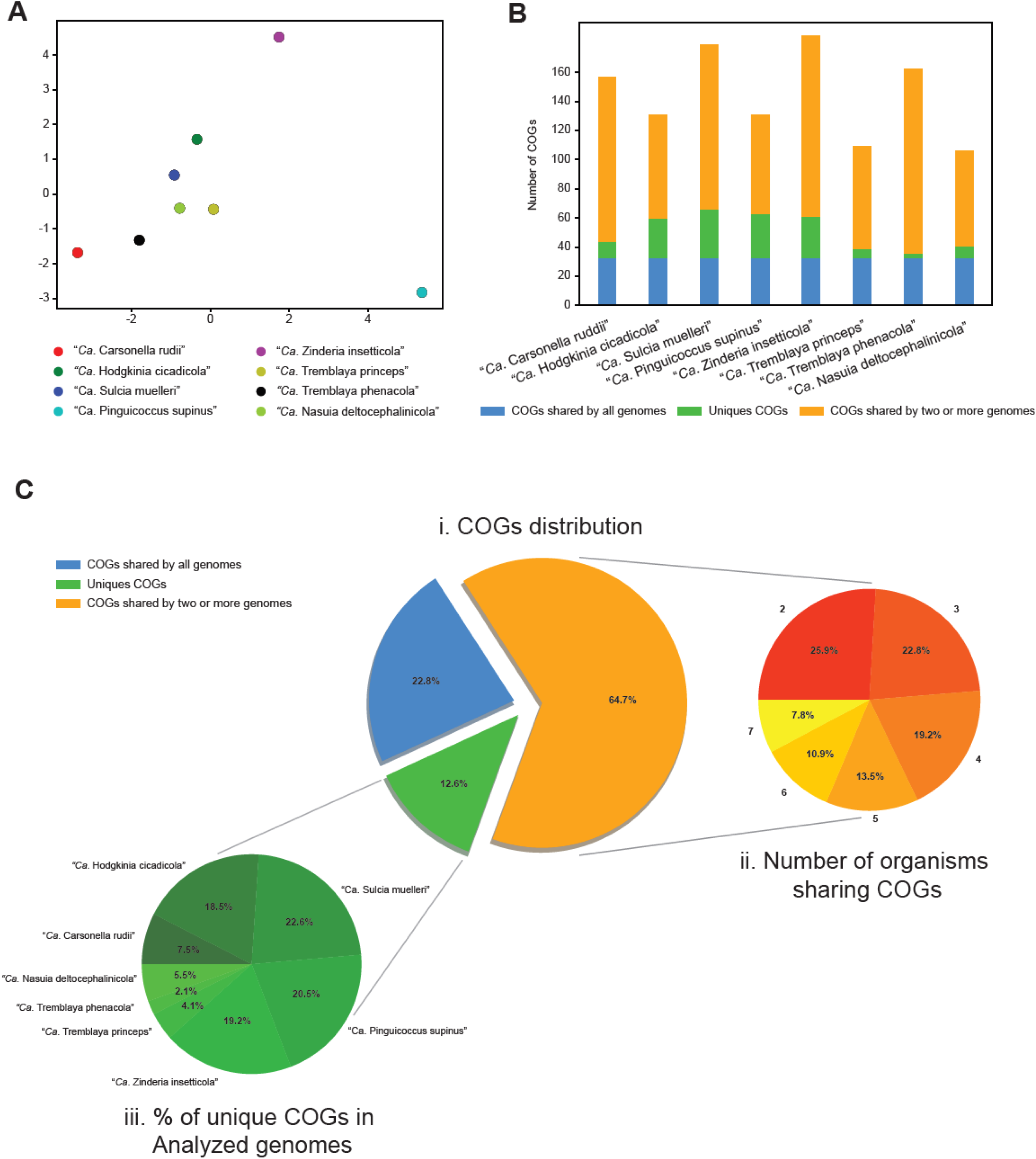
COG analysis of “*Candidatus* Pinguicoccus supinus” and other bacteria with highly reduced genome. A) Principal Component Analysis of numerosity in COG classes; explained variance by Component one 32%; explained variance by Component two 24%; B) Distribution of COGs in each analysed genome; C) Distribution of COGs, showing: i. percentage of COGs shared by all the analysed genomes, COGs unique for each genome, and COGs shared by at least two genomes; ii. Number of organisms sharing COGs. Each set groups together the COGs shared by a given number of organisms (i.e. 1, 2, 3…7), regardless of their identity; iii. Percentage of unique COGs in analyzed genomes.

While almost all the bacteria with highly reduced genome cluster together (Fig. 6a), the variance explained by the first component positions “*Ca.* Pinguicoccus supinus” remarkably far from them. This component is mainly correlated with the COG classes Q - Secondary metabolites biosynthesis, transport and catabolism (0.36), I - Lipid transport and metabolism (0.34), and K - Transcription (0.33). The variance explained by the second component separates “*Ca.* Zinderia insecticola” from the other bacteria with highly reduced genome. This component is mainly correlated with COG classes P - Inorganic ion transport and metabolism (0.42), H - Coenzyme transport and metabolism (0.39), and C - Energy production and conversion (0.33) classes.

The COG analysis also showed the presence of a set of 33 genes shared by all the small genome bacteria in analysis (Figs 6b, c), mostly related to DNA replication, transcription, and translation. Although the metabolic capability of “*Ca.* Pinguicoccus supinus” is, in general, similar to that of other highly reduced genomes (88% of COGs shared with at least another analysed genome; Figs 6b, c) (McCutcheon et al. 2012; Moran and Bennett 2014), the novel genome lacks the capability to synthesize any amino acids. Moreover, no catalytic subunit of the DNA polymerase was identified. “*Ca.* Pinguicoccus supinus” possesses 30 genes (12% of the total retrieved COGs in this bacterial genome) that are absent in all the other tiny genomes, mostly related to COG classes I (lipid transport and metabolism) and M (cell wall/membrane/envelope biogenesis). In general, with respect to the other analyzed bacteria, this endosymbiont includes a richer set of genes involved in fatty acid biosynthesis, in glycosylation and glycan modification (Supplementary Table 8).

#### Phylogeny and phylogenomics

The 16S rRNA gene-based phylogeny showed “*Ca.* Pinguicoccus supinus” as a member of the family *Puniceicoccaceae* (*Verrucomicrobia, Opitutae*) (Fig. 7). It clustered with sequences from uncultured organisms, forming a clade related to the genera *Coraliomargarita, “Fucophilus”, Cerasicoccus*, and *Ruficoccus* (Fig. 7). The long branch of “*Ca.* Pinguicoccus supinus” suggests a higher evolutionary rate with respect to related *Verrucomicrobia*. Moreover, we observed that the inclusion of the “*Ca.* Pinguicoccus supinus” sequence in the *Verrucomicrobia* tree reduces the values of statistical supports for the nodes of the *Puniceicoccaceae* clade (data not shown). This is consistent with the occurrence of a long branch attraction phenomenon between “*Ca.* Pinguicoccus supinus” and the outgroup, which destabilizes *Puniceicoccaceae* despite high taxon sampling (Bergsten 2005).

**Figure 7.**
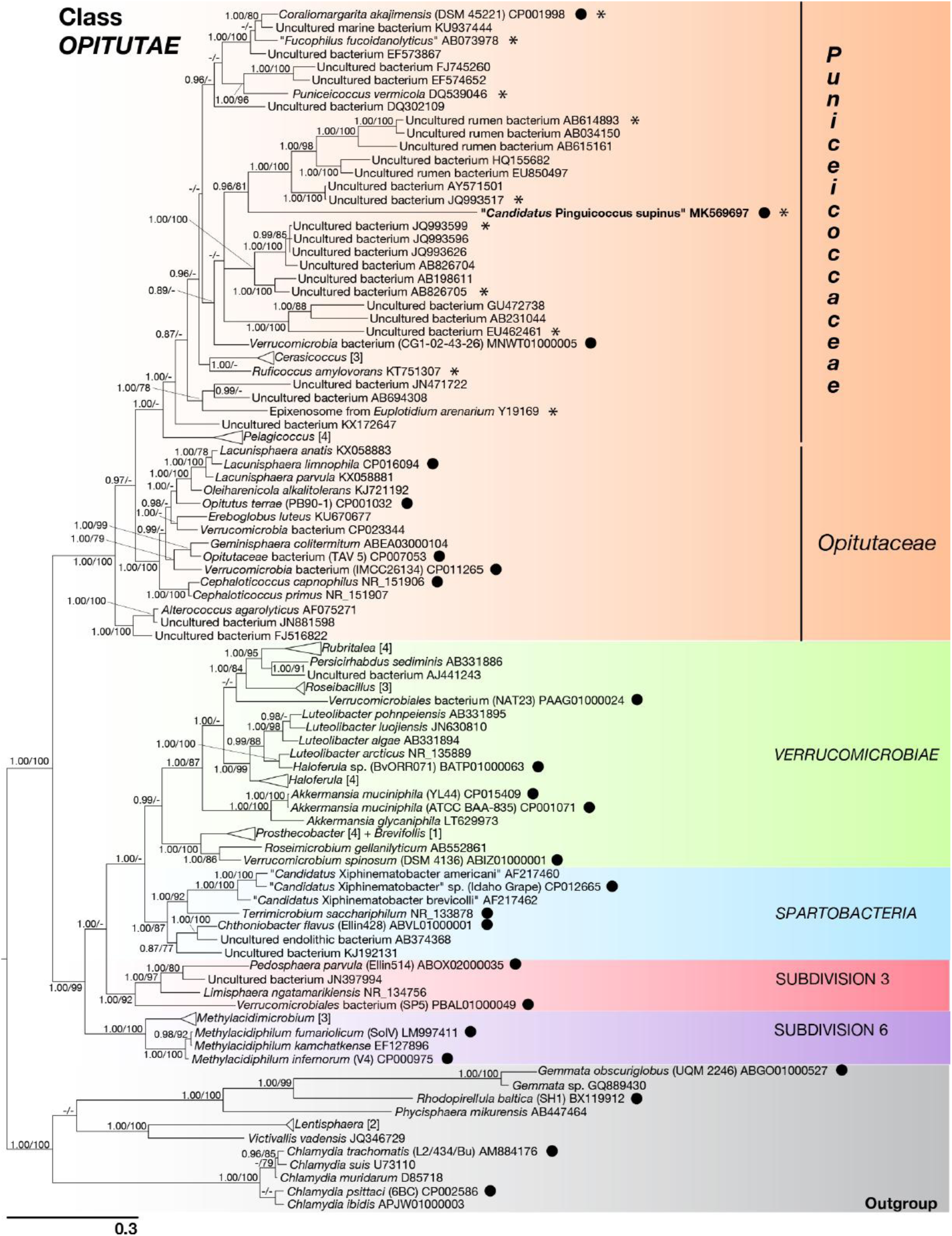
Phylogenetic tree of Phylum *Verrucomicrobia*, based on the 16S rRNA gene. The phylogenetic position of “*Candidatus* Pinguicoccus supinus” is shown. Numbers associated to nodes represent posterior probability and bootstrap values, respectively (only values above 0.80–75 are shown). Black circles indicate organisms also employed in phylogenomic analysis (Fig. 8). Asterisks indicate sequences employed in the 16S rRNA gene screening on IMNGS (Fig. 10; Supplementary Table 9). Numbers in square brackets, associated to collapsed branches, indicate how many sequences are not shown (for list of hidden sequences see Supplementary Table 4). Sequence obtained in the present work is in *bold*.

The result of the phylogenomic analysis is fully consistent with the phylogeny of the 16S rRNA gene, confirming the position of the endosymbiont inside the phylum *Verrucomicrobia* and in the class *Opitutae* (Fig. 8). “*Ca.* Pinguicoccus supinus” clustered together with uncultured organisms (GB_GCA_001872735, GB_GCA_002309885, GB_GCA_002336265), in a clade related to *Coraliomargarita akajimensis* (RS_GCF_000025905), and thus inside the family *Puniceicoccaceae* (Fig. 8).

**Figure 8.**
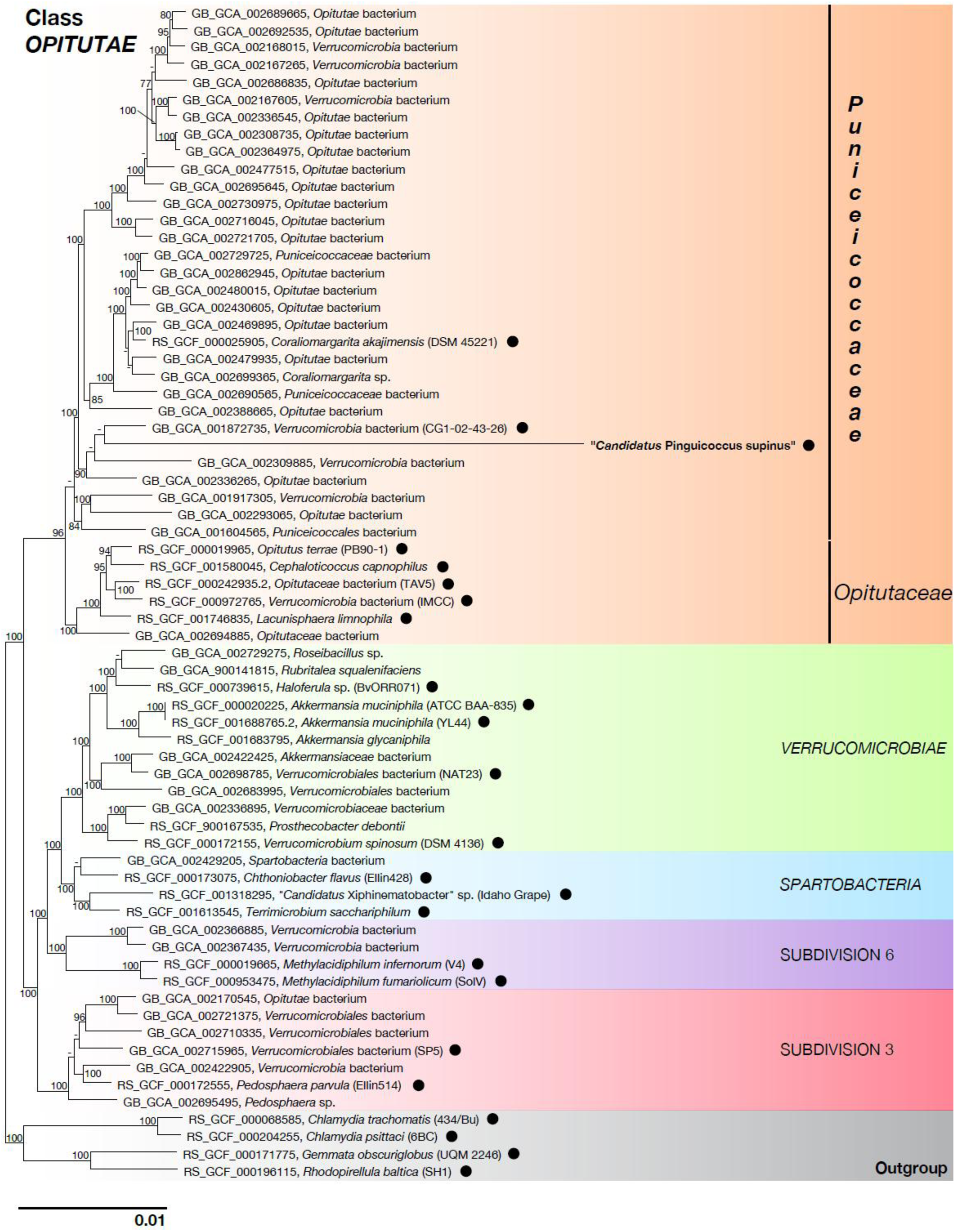
Phylogenomic tree of *Verrucomicrobia*, showing evolutionary relationships of “*Candidatus* Pinguicoccus supinus”. Numbers associated to nodes represent bootstrap values (only values above 75 are shown). Black circles indicate organisms also employed in phylogenetic analysis (Fig. 7). Genome of “*Ca.* Pinguicoccus supinus” obtained in the present work is in *bold* (accession number: CP039370).

#### Localization inside host cell

FISH experiments showed 100% of *E. vanleeuwenhoeki* cells positive to specifically designed probes for “*Ca.* Pinguicoccus supinus” (Fig. 9). The number of endosymbionts per host cell ranged from 12 to 36 bacteria (*X*±SD: 25.1 ± 7.3; n=16). The symbionts showed a peculiar pattern of distribution in almost all the observed *Euplotes* cells: beneath the cortex of the host, in clusters of several bacteria grouped at one side of the cell (Fig. 9c), or more commonly at the two poles (Fig. 9f). These observations support data from TEM analysis, showing “*Ca.* Pinguicoccus supinus” generally lying close to the cortex and forming clusters with other conspecifics.

**Figure 9.**
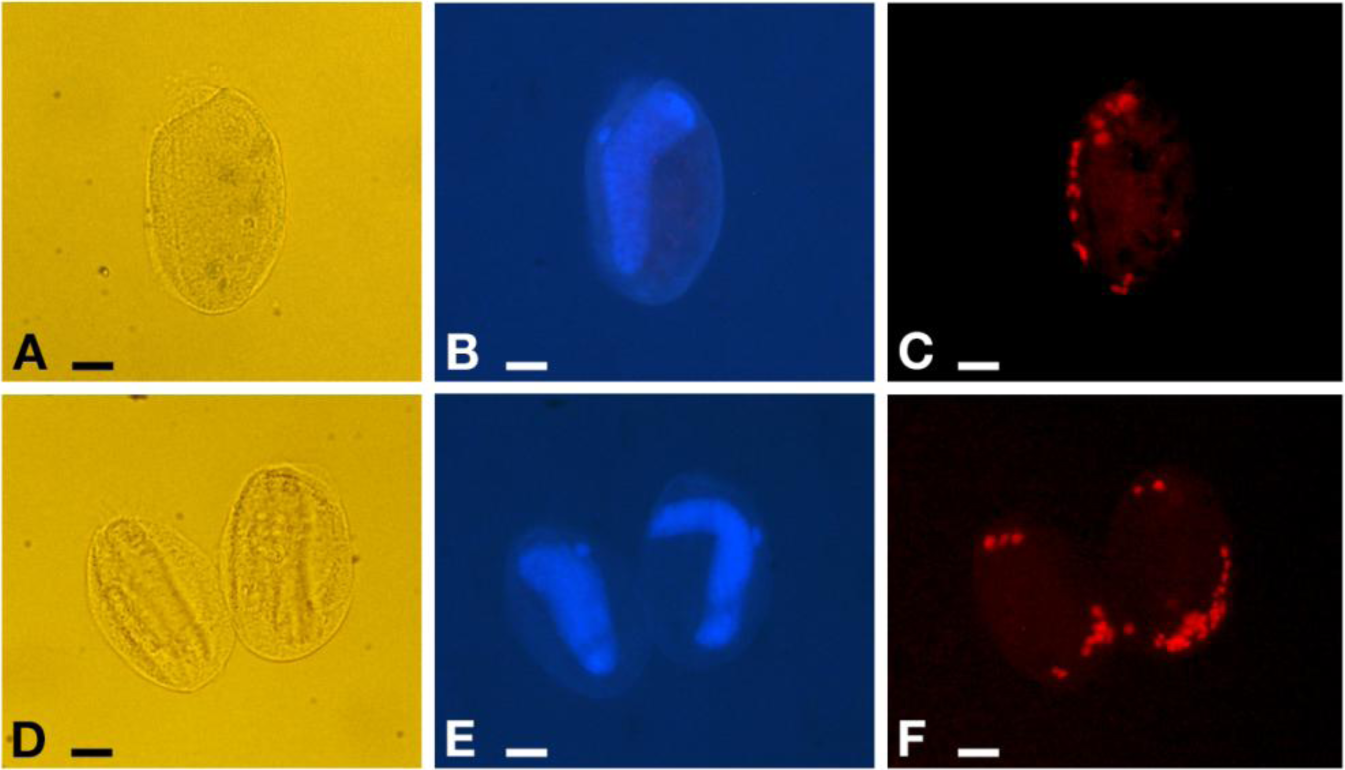
Fluorescence *in situ* hybridization experiments on *Euplotes vanleeuwenhoeki* sp. nov. A-C) Specimen hybridized with probe EUB338 VII. D-F) Two specimens hybridized with probe Pingui_1174. A), D) Pictures at DIC microscope of fixed cells; B), E) DAPI staining, showing position of nuclear apparatus; C) Cell positive to probe EUB338 VII, fluorophore emitting in red (Cyanine-3); F) Cell positive to probe Pingui_1174, fluorophore emitting in red (Cy-3). *Bars* stand for 10 µm.

#### Diversity and environmental screening of the endosymbiont

The IMNGS platform was used to investigate the molecular diversity and environmental distribution of sequences showing high identity with the novel endosymbiont of *Euplotes*. The 16S rRNA gene of the endosymbiont was queried at 95% similarity threshold, and a total number of 4,572 sequences were obtained and subsequently used to investigate the molecular diversity of 16S rRNA gene hypervariable regions. Sequences were clustered in 90 OTUs with 99% threshold identity, then OTUs were grouped according to their hypervariable region in V1-V2, V4-V6 and V7-V8, and phylogenetic trees were inferred accordingly (Fig. 10). Phylogenetic trees showed that six diverse groups can be distinguished for all hypervariable regions considered, and their phylogenetic relations are in agreement with those shown by the full-length 16S rRNA gene phylogenetic reconstruction (Fig. 10). The first group (red) includes the novel endosymbiont and related sequences (AB614893, JQ993517; Fig. 7), the second one (violet) is formed by sequences from uncultured bacteria (JQ993599, AB826705), the third one (purple) consists of a single uncultured bacterium (EU462461), the fourth one (green) includes *Coraliomargarita akajimensis* (CP001998), “Fucophilus fucodainolyticus” (AB073978), and *Puniceicoccus vermicola* (DQ539046), the fifth one (yellow) has *Cerasicoccus frondis* (AB372850) and *Ruficoccus amylovorans* (KT751307), and the last one (blue) includes just epixenosomes from *Euplotidium arenarium* (Y19169) (Fig. 10). The IMNGS environmental screening also showed that metagenomic samples positive to “*Ca*. Pinguicoccus” were very limited in number, namely 0.01% of total samples. Indeed, the related sequences were present in just 32 out of a total of 303,362 samples (Supplementary Table 9). The environmental origin of positive samples is reported in Supplementary Table 9. Positive hits originated from very diverse samples, such as seawater, wastewater, microbial mats, but also potential host organisms, such as shrimps and plants (Supplementary Table 9). However, the overall abundance of positive hits was extremely low and always below 1% within each sample. The highest abundance was found in samples originating from a study on wastewater (Al-Jassim et al. 2015), where “*Ca*. Pinguicoccus” abundance ranged between 0.005 and 0.752% (Supplementary Table 9).

**Figure 10.**
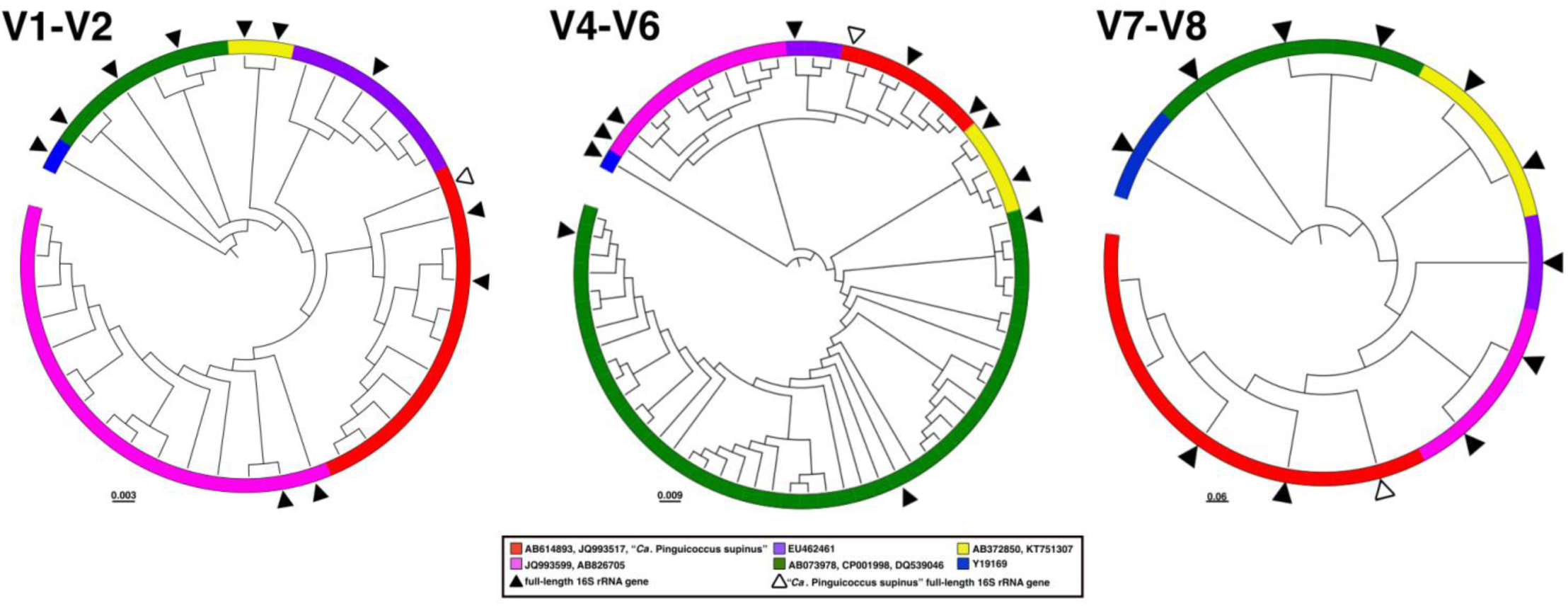
Diversity of “*Candidatus* Pinguicoccus supinus” based on 16S rRNA gene amplicon search in IMNGS. 16S rRNA gene hypervariable regions phylogenetic trees. OTUs were clustered with 99% identity and were longer than 300 bp. A total number of 90 OTUs were separated for each hypervariable region taken into analysis: 29 for V1-V2, 60 for V4-V6, 2 for V7-V8. Complete 16S rRNA gene was employed in the analysis to enlighten the diversity of “*Ca*. Pinguicoccus supinus”.

## Discussion

### Next generation taxonomy

The present work aims to introduce and describe an innovative approach to the characterization of living beings, integrating the holobiont concept into traditional and integrative taxonomy leveraging the power of high-throughput DNA sequencing. We propose for this approach the name “next generation taxonomy”.

To exemplify the concept of “next generation taxonomy” applied to holobionts, we used the ciliate *E. vanleeuwenhoeki* sp. nov. and its symbiont, “*Ca.* Pinguicoccus supinus” gen. nov., sp. nov., as a case study. In the following discussion, we propose an updated pipeline for the description of holobionts starting with the sequential description of each biont of the system.

### First biont: Euplotes vanleeuwenhoeki

#### Comparison with congeners

*Euplotes vanleeuwenhoeki* showed morphological and molecular affinity with other members of the genus, such as *E. antarcticus* (Fenchel and Lee 1972; Petz et al. 1995)*, E. trisulcatus* (Carter 1972)*, E. euryhalinus* (Valbonesi and Luporini 1990a)*, E. charon* (Song and Packroff 1997; Shao et al. 2010), and *E. magnicirratus* (Carter 1972). Indeed, they share the same dargyrome and frontoventral cirri patterns (for details see Table 2), and clustered together in our molecular phylogeny (Fig. 3). As for its morphology, *E. vanleeuwenhoeki* was particularly similar to *E. trisulcatus* (Carter 1972). Nevertheless, the combination of several characters (i.e. posterior end rounded *versus* pointed; peristome length, number of AZM, number of dikinetids in mid-dorsal row - for details see Table 2), the overall body shape, the type of habitat (freshwater *vs* marine), and the 18S rRNA gene sequence supported its attribution to a novel species.

**Table 2.**
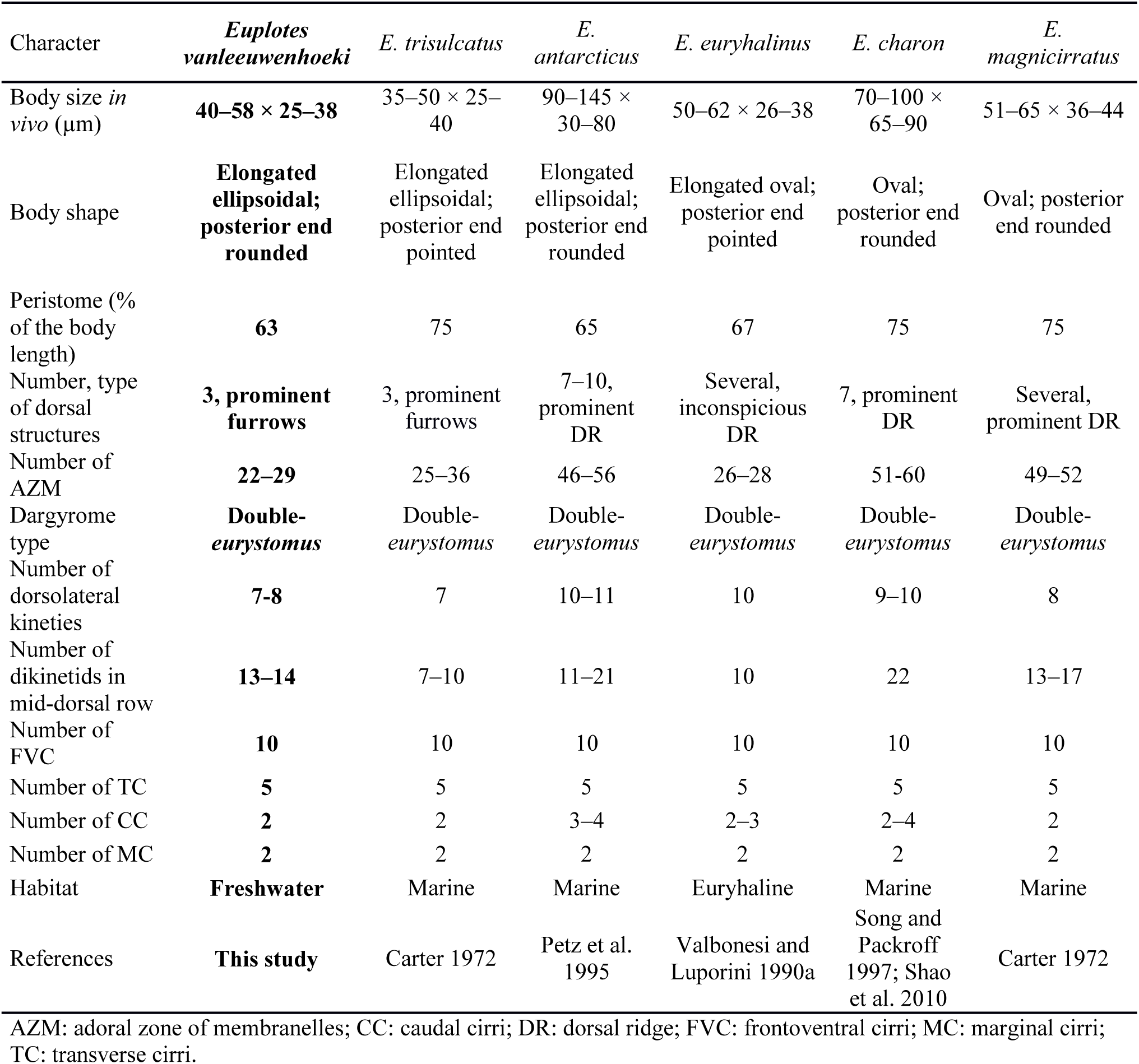
Morphological comparison between *Euplotes vanleeuwenhoeki* sp. nov. and selected congeners

Unfortunately, the sequences closest to that of *E. vanleeuwenhoeki* in our phylogeny reconstruction (i.e. *E. magnicirratus* – AJ549210 (Petroni et al. 2002); *E. charon* - AF492705 (Li and Song 2006); *E. euryhalinus* – EF094968 (Vallesi et al. 2008), JF903799 (Keerthi and Reddy unpublished); *E. trisulcatus* EF690810 (Schwarz and Stoeck unpublished), *Euplotes* cf. *antarcticus* – FJ998023 (Gao and Song unpublished) are not accompanied by any morphological description. Consequently, data used in our morphometric comparison inevitably derive from previous, exclusively morphological, studies (i.e. Carter 1972; Petz et al. 1995; Valbonesi and Luporini 1990a; Song and Packroff 1997; Shao et al. 2010). Moreover, there is no certainty that above-mentioned *Euplotes*-derived sequences have been properly attributed to the correct ciliate species; thus, a certain cautiousness is recommended until a comprehensive redescription of such organisms, joining morphology and molecular data, is performed. In this somehow unclear landscape, for the sake of completeness, we decided to include in our comparison among different *Euplotes* species (Table 2) also *E. charon*, because it shares some basic features with *E. vanleeuwenhoeki*, (i.e. dargyrome, and frontoventral cirri pattern), although the phylogenetic position of this species is far from being resolved. Indeed, the sequence AF492705, under the name of *E. charon*, is not linked to any morphological description (Li and Song 2006); this sequence was used by Shao and colleagues (Shao et al. 2010) as molecular reference in their redescription of *E. charon*, for which, unfortunately, they used a different *Euplotes* strain for morphological analysis. However, there are several other available sequences that are named after *E. charon*, but they all are not supported by any morphological data (i.e. AJ305249, Petroni et al. 2002; JF694043, Huang et al. 2012; FJ87077-80, Huang and Song unpublished) and these sequences cluster far away from *E. vanleeuwenhoeki.* A critical revision of the species *E. charon* using at least integrative taxonomy is consequently highly recommended.

As a general remark, *E. vanleeuwenhoeki* share with many other species the plesiomorphic condition of the genus, i.e. the occurrence of 10 FVC and a double dargyrome of *eurystomus*-type (e.g.: *E. alatus, E. antarcticus, E. balteatus, E. charon, E. crenosus, E. curdsi, E. enigma, E. euryhalinus, E. focardii, E. harpa, E. inkystans, E. magnicirratus, E. neapolitanus, E. octocirratus, E. palustris, E. parabalteatus, E. platysoma, E. plicatum, E. polycarinatus, E. qatarensis, E. quinquecarinatus, E. rariseta, E. trisulcatus*, and *E. tuyraui*). Although these two features are always reported in morphological description, due to their plesiomorphic status they are of limited use for taxonomic identification. As an example, some authors (Jones and Gates 1994) even proposed to synonymise almost all species possessing 10 FVC and a double dargyrome under the name of *E. charon*. Obviously, nowadays, this proposal is not acceptable on the basis of molecular data (Boscaro et al. 2018; Lian et al. 2018) and because of other morphological features characterizing some of the organisms (e.g. dorsal furrows/ridges, and body shape and proportions). It has been proved that, in many cases, this ancestral condition underwent severe, independent modifications during the evolutionary history of the genus (Petroni et al. 2002; Syberg-Olsen et al. 2016). Indeed, the double dargyrome evolved in other different patterns, such as “single”, “double *patella*-type”, “multiple” (Carter 1972), and “complex”, in many distinctive events (Syberg-Olsen et al. 2016). The dargyrome modification was explained either by increasing in number or by unification of alveoli (Valbonesi and Luporini 1995; Syberg-Olsen et al. 2016). It is not by chance that some *Euplotes* species present a great phenotypic plasticity, determined by particular physiological and/or ecological conditions, such as the presence of a predator and/or food availability (i.e. *E. focardii* (Valbonesi and Luporini 1990b), *E. balteatus* (Tuffrau 1964), *E. octocarinatus* (Kuhlmann and Heckmann 1994; Kopp and Tollrian 2003), and *E. variabilis* (Tuffrau et al. 2000)). Similarly, one or more FVC were independently lost many times, as also evidenced by the vestiges still detectable in some *Euplotes* species (Washburn and Borror 1972; Petroni et al. 2002; Jiang et al. 2010; Syberg-Olsen et al. 2016).

#### Mitochondrial genome

Ciliates possess very peculiar mitochondrial genomes, being among the first to be identified as linear (Suyama and Miura 1968; Morin and Cech 1986) and showing several split rRNA genes (Schnare et al. 1986) and protein genes (Burger et al. 2000). To our best knowledge, in ciliates, the set of potentially split genes in the mitochondrial genome includes *nad1, nad2, rps3, rnl* and *rns*. This list now includes the *cox1* gene, which is split in *E. vanleeuwenhoeki*. Considering that the splits in *nad1, nad2, rps3, rnl* and *rns* genes occur approximately at the same positions in the mitochondrial genomes of Spirotrichea, it has been hypothesized they could be present in the last common ancestor of *Euplotes* and *Oxytricha* (Swart et al. 2011). However, since the *cox1* gene is split only in the *E. vanleeuwenhoeki* genome, it is likely that this represents a more recent event. The occurrence of multiple and evolutionary independent split genes in ciliates could be considered indicative of a possible inherently higher tolerance to split events in mitochondrial genes in these organisms. The mitochondrial genome of *E. vanleeuwenhoeki* shows a high level of colinearity with the other two available *Euplotes* mitochondrial genomes. In addition, all *Euplotes* share a good colinearity with *O. trifallax* (two single-gene inversions and one single-gene transposition). Since the amino acid substitution rates in ciliate mitochondrial genomes appear to be high (Swart et al. 2011), this result enforces the idea that mitochondrial genome colinearity can be employed in phylogenetic analysis as a valid supporting feature to define the systematics of ciliates, or to distinguish different taxonomic levels among very related species using the non-colinear terminal regions (Blanchette et al. 1996; Boore and Brown 1998). Indeed, mitochondrial genes, such as *cox1*, *cox2* and *rns*, are being increasingly used in taxonomy, both for phylogenetic analysis (Dumilag et al 2018), and for discriminating between species with similar morphological features. For example, mtDNA genes have been employed to reveal several cryptic species in Ciliophora (Barth et al. 2006; Doerder 2019) and metazoan such as annelids (Nygren et al 2018) and insects (Ya’cob et al 2017), or to discriminate between subspecies of the bee *Apis mellifera* (Ilyasov et al 2016). The limited number of available mitochondrial genome sequences of ciliates currently does not allow to perform phylogenetic analyses based neither on sequences nor on colinearity. Nevertheless, we predict that such analyses will be extremely helpful, in the future, to resolve phylogenetic and taxonomic issues as already occurred in other taxa (e.g. metazoa) in which more sequences are already available (Lavrov et al. 2004; Cameron 2014). We expect the same for ciliates as soon as a sufficient number of novel mitochondrial genome sequences will be available.

#### Symbiosis with a Verrucomicrobia member

Members of genus *Euplotes* are known to host symbiotic eukaryotes, mutualistic such as endocellular algae (Diller and Kounaris 1966; Lobban et al. 2005), or opportunistic such as microsporidia (Fokin et al. 2008) and trypanosomes (Fokin et al. 2014). Moreover, *Euplotes* species display a certain attitude to harbor one (Heckmann et al. 1983; Vannini et al. 2004, 2013), or even multiple bacterial symbionts, this second condition being more common if the ciliate presents the obligate symbiosis with the betaproteobacterium *Polynucleobacter necessarius* (Heckmann and Schmidt 1987; Boscaro et al. 2012, 2013b; Schrallhammer et al. 2013; Vannini et al. 2010, 2014; Boscaro et al. 2018; Chiellini et al. 2019). In those cases, the secondary or tertiary symbionts, if present, frequently belong to *Alphaproteobacteria*, e.g. *Rickettsiaceae* (Schrallhammer et al. 2013; Vannini et al. 2014, Lanzoni et al., 2019) or “*Ca*. Midichloriaceae” (Vannini et al. 2010; Boscaro et al., 2013b; Senra et al. 2015). Also, *Gammaproteobacteria* were retrieved in different *Euplotes* species (Schrallhammer et al. 2011; Boscaro et al. 2012; Chiellini et al. 2019; Vallesi et al. 2019).

To date, “*Ca*. Pinguicoccus supinus” as endosymbiont of *E. vanleeuwenhoeki* constitutes a *unicum*, being the first member of *Verrucomicrobia* with a highly reduced genome, and also the first found as endosymbiont of a protist. Indeed, bacteria of the phylum *Verrucomicrobia* are mostly free-living organisms, nearly ubiquitous, present both in the soil and in wet environments (Zhang and Xu 2008; Freitas et al. 2012), and often retrieved in extreme habitats (Pearce et al. 2003; Hou et al. 2008). Little is known about *Verrucomicrobia* living as obligate symbionts, which at present were described in nematodes (Vandekerckhove et al. 2000), echinoderms (Sakai et al. 2003), squids (Collins et al. 2012), and in gut of different metazoans (Yildirim et al. 2010; Romero-Pérez et al. 2011), including humans (Wang et al. 2005). The only other case of association of *Verrucomicrobia* with protists is the symbiosis between the ciliate *Euplotidium* and the so-called epixenosomes (Rosati et al. 1998; Petroni et al. 2000; Modeo et al. 2013b), which are verrucomicrobial ectosymbionts able to extrude and defend their host against predators (Rosati et al. 1999).

### Second biont: *“Candidatus* Pinguicoccus supinus”

#### Endosymbiont morphology

The morphology of some PVC members is renown to be peculiar. Indeed, some of them present a certain degree of cellular compartmentalization, even if some aspects of the subject are still under debate (Lindsay et al. 2001; Fuerst and Sagulenko 2011; Devos 2014). In particular, cell compartmentalization has been proposed for some members of *Verrucomicrobia*, such as *Prostechobacter dejongeii* (*Verrucomicrobiaceae*) and *Coraliomargarita akajimensis* (*Puniceicoccaceae*) based on molecular and ultrastructural studies (Lee et al. 2009; Pinos et al. 2016). In those two bacteria, the presence of two different cellular compartments, delimited by an intracytoplasmic membrane, has been described. These two compartments were recognizable as a ribosome-free region, the paryphoplasma, and a region containing ribosomes and nucleoid, the pirellulosome. In this framework, some morphological traits of “*Ca.* Pinguicoccus supinus”, such as membrane invaginations and folding, could be considered homologous to those observed in other *Verrucomicrobia* and PVC members in general, although our data, based solely on TEM observation, definitely do not provide clear evidence of cell compartmentalization. A possible future development could be the use of cryo-electron tomography (Medeiros et al. 2018).

#### Endosymbiont genome

The genome of “*Ca*. Pinguicoccus supinus” was found to be extremely small (163,218 Kbp), being, to our best knowledge, the fourth smallest bacterial genome sequenced to date (smallest being “*Ca.* Nasuia deltocephalinicola”: 112,031 bp) (Bennet and Moran 2013). The novel genome is highly reduced and lacks many genes, especially compared with the relative free living *Verrucomicrobia* (range 2.2 – 7.3 Mbp; Hou et al. 2008; Kotak et al. 2015) but also with other *Verrucomicrobia* symbionts, such as the ones belonging to the genus “*Ca*.

Xiphinematobacter” (~916 Kbp) (Brown et al. 2015). The tiniest genomes (<500 Kb) are found in putatively ancient mutualistic symbioses, where symbionts are beneficial to their hosts’ metabolism and hosts have in turn evolved to support and control the symbiosis (McCutcheon and Moran 2012; Bennet and Moran 2013; Moran and Bennett 2014; Kobiałka et al. 2016). Such levels of co-evolution often also imply phenomena of severe genome reduction, with gene loss and often a subsequent modification of the nucleotide base composition (Moran and Bennet 2014). This framework gives us some clues to try to understand the peculiar characteristic of the novel symbiont “*Ca*. Pinguicoccus supinus”. All other symbionts with highly reduced genomes provide their hosts with nutrients that are absent in their diet. In those cases, the nutritional support role is clear, due to the fact that they retain some metabolic pathways for the biosynthesis of amino acids essential for their hosts (Wilson et al. 2010; for a review see Moran and Bennet 2014). Moreover, several of these symbionts are co-obligate with other bacteria complementing each other for the enzymatic repertoire necessary to provide nutritional support to the host. This seems not to be the case of “*Ca*. Pinguicoccus supinus”, which is devoid of genes related to amino acids, co-factors or vitamins biosynthesis, and is the only symbiont retrieved in the *E. vanleeuwenhoeki* holobiont. The extensive gene loss and genome modification suggest that this symbiosis may have an ancient origin, and that it may play an important role for the host, although its function is yet to be understood. This case could be partly similar to that of the well-ascertained mutualist bacterium, *Polynucleobacter necessarius* in other *Euplotes* spp.. Indeed, this bacterium is necessary for the survival of its hosts, but no precise clues on the metabolic nature of the interaction have been provided by genomic analyses (Boscaro et al. 2013c, 2018). This has been linked to the peculiar ecology of ciliates, which normally feed on free-living bacteria, and are thus unlikely to require dietary compensations from bacterial endosymbionts. It has been hypothesized that they may require help in catabolism or other undefined functions, and such a contribution would not be simple to detect by analyses of the symbionts’ genomes only (Boscaro et al. 2013c).

The comparison between “*Ca.* Pinguicoccus supinus” and the other symbionts with highly reduced genomes shows the presence of a core set of 33 genes retained by all these evolutionary unrelated bacteria. Those genes are mostly involved in DNA replication, transcription, and translation, as well as in protein folding and stability. Clearly, these core cellular functions are required even in bacteria with such extremely reduced genomes. Interestingly, in “*Ca.* Pinguicoccus supinus”, it was not possible to find a gene homologous to DNA polymerase II catalytic subunit (*dnaE*), or, in general, to any other protein with predicted DNA polymerase catalytic activity. This is a striking difference with respect to the other analysed symbionts, and, to our best knowledge, it is a unique case among all bacteria. A Blast analysis on the entire preliminary assembly of the whole holobiont did not reveal the presence of any *dnaE* gene, suggesting that there is no such gene integrated in the host genome. This finding strongly suggests that “*Ca.* Pinguicoccus supinus” relies on proteins obtained from the host to replicate its own genome, analogously to what has been proposed for other essential cellular functions for other symbionts with highly reduced genomes (Moran and Bennet 2014).

A common feature of the previously characterized symbiotic bacteria with highly reduced genomes is the complete lack of genes related to the production of components of the cellular envelope such as phospholipids, lipopolysaccharide, peptidoglycan and related membrane proteins (Moran and Bennett 2014). Unique in this regard, “*Ca.* Pinguicoccus supinus” presents the pathway for the initiation and elongation of fatty acids, and several glycosyl transferases, although the phospholipid synthesis pathway is missing. Obligate small genome symbionts of insects are found in specialized cells, the bacteriocytes and some symbionts of unicellular eukaryotes are located in a specific cellular structure, the symbiosome, while “*Ca*. Pinguicoccus supinus” resides free in the host cytoplasm. This could imply the requirement for a more complex regulation of the membrane structure (e.g. variation of fatty acids length in phospholipids) to face a “less stable” environment, thus possibly explaining the unique genes found in the genome of “*Ca*. Pinguicoccus supinus”. Interestingly, “*Ca.* Pinguicoccus supinus” has often been observed in close contact with host’s lipid droplets. This not yet elucidated interaction with host’s lipids could be another reason for the retaining of genes devoted to fatty acid biosynthesis.

The genome of “*Ca.* Pinguicoccus supinus” presents a non-standard genetic code (NCBI genetic code “4”), in which the canonical stop triplet UGA is recoded for tryptophan, consistently with the absence of release factor 2, implied in the normal recognition of UGA as stop. This is consistent with other highly reduced genomes (McCutcheon et al. 2009) and has been tentatively linked to a directional mutation pressure related to the AT/GC composition (Osawa and Jukes 1988), or directly linked to genome reduction and energetic constrains (McCutcheon et al. 2009).

#### Endosymbiont phylogeny

Both the phylogenetic and phylogenomic analyses, the first including more taxa and the second more genes, support the position of “*Ca.* Pinguicoccus supinus” in the family *Puniceicoccaceae* (*Opitutae*, *Verrucomicrobia*). We suggest that the long branch of the novel species is due to an overall high evolutionary rate, and consequently may decrease support for some nodes inside the family-clade.

It is worth to notice that our phylogenetic and phylogenomic analyses are among of the most taxonomically exhaustive up to date among *Verrucomicrobia* and resulted coherent with those from previous studies (Yoon et al. 2010; Yoon 2011; Kim et al. 2015; Parks et al. 2018). They also highlight the necessity of systematic revision for some species and taxa (i.e. *Brevifollis gellanilyticus* clustering within genus *Prosthecobacter*; Subdivision 3 and Subdivision 6 still waiting for a proper naming).

#### Endosymbiont diversity and environmental distribution

In order to evaluate how widespread and abundant the novel taxon is, we evaluated the presence of sequences related to 16S rRNA gene of *E. vanleeuwenhoeki* endosymbiont in IMNGS, which resulted to be very limited. The main possible reason of the scarce abundance of “*Ca*. Pinguicoccus supinus” and related sequences in online repositories resides in the fact that 16S rRNA gene primers employed in metabarcoding and metagenomic studies often do not detect certain bacterial groups, such as *Verrucomicrobia* (Bergmann et al. 2011; Takahashi et al. 2014). Indeed, the choice of primers used in such NGS analysis should be carefully pondered as some bacterial taxa might be undetected, thus not depicting the real diversity in an environment. Our analysis shows that the 16S rRNA sequences related to the endosymbiont were present in just 0.01% of total samples, and never reached 1% abundance in positive samples, a number of which resulted to be microbiome studies of multicellular organisms (Supplementary Table 9). The hypervariable region diversity analysis confirmed the limited presence of sequences from these bacteria sequence in metabarcoding samples. Extracting from the databases sequences similar to “*Ca.* Pinguicoccus supinus” and performing phylogenies on the hypervariable regions of the 16S rRNA gene, six diverse groups could be identified, and their phylogenetic relationships retraced those of the complete 16S rRNA gene phylogeny (Figs 7, 10). Moreover, hypervariable region diversity analysis pointed out that “*Ca*. Pinguicoccus supinus” was usually a stand-alone sequence within its group, with the only exception of V4-V6 phylogenetic tree having a single OTU associated to the endosymbiont sequence (Fig. 10). On the contrary, the other related *Puniceicoccaceae*, which were isolated and characterized from the environment (mostly marine), clustered in groups encompassing numerous OTUs (Fig. 10). The same stand-alone phenomenon occurred also for the epixenosomes, the aforementioned ectosymbionts of the ciliate *Euploditium arenarium*, that always formed a defined and separated lineage in all our analyses (Fig. 10). The limited presence of OTUs associated to these microorganisms could be also explained with the symbiotic nature of these bacteria, which occupy a peculiar and circumscribed ecological niche, possibly making their detection more complicated. It is also worth to notice that the number of OTUs *per* clade is significantly different when region V1-V2 is compared to region V4-V6 (Fig. 10). Indeed, in region V1-V2 “*Ca*. Pinguicoccus supinus” and related clades (in red) are relatively richer in OTUs if compared to region V4-V6, whereas the opposite is true for the *Coraliomargarita*-“*Fucophilus*”-*Puniceicoccus* clade (in green). This observation suggests that commonly used primers for region V1-V2 and V4-V6 could recognize the two clades with a different efficiency, producing artefactual results in terms of abundance and biodiversity.

### The holobiont paradigm as a possible new standard for the “next generation taxonomy” of unicellular organisms

The concept that “host plus symbionts” constitute a single functional unit, with its own characteristics, different from and new with respect to those of the specific parts, is nowadays accepted by many researchers, albeit with different interpretations (Rosenberg et al. 2007; Bordenstein and Theis 2015; Bosch and Miller 2016; Roughgarden et al. 2018).

Therefore, to our opinion, it seems opportune, if not even necessary, to treat the sum of the different parts of this system in a unitary and integrated way, starting from the very beginning, i.e. the taxonomic description. Hence, we propose the idea of shifting attention of taxonomy from unicellular host and/or single symbionts to the whole holobiont. This approach integrates all the associated species characterizations into the holobiont description, providing not only morphological characters and molecular markers, but also functional and associative parameters.

According to this line of thought, such a multidisciplinary approach represents a considerable improvement, allowed by the development of more sophisticated and powerful tools in molecular biology and genomics. In details, here we suggest the inclusion of two additional descriptive parameters: the host’s mitogenome and the characterization of the possibly present associated bionts, integrating novel useful features to the description of organisms. This does not imply that the traditional taxonomic tools are to be considered obsolete, and they will indeed continue to be the basis for species descriptions.

A question naturally arising is what should be considered a “holobiont”. The system constituted by the unicellular host plus its bacterial symbiont(s) perfectly fits with the definition of “holobiont”. As we pointed out earlier in the Introduction section, we suggest treating as a single holobiont the compound of different organisms in stable associations, which we can consider to be forming a single integrated functional and ecological unit. These systems fit the definition of symbiosis *sensu* de Bary, i.e. including parasitism. Indeed, in studies on symbiosis, a long lasting, yet unresolved, debate is how to define the boundaries among different kinds of symbiosis (e.g. ranging from mutualism to parasitism) (Goff 1982; Martin and Schwab 2012). The threshold could be particularly blurred for interactions between protists and other unicellular organisms. In practice, this “gray area” in between is frequently due to the difficulty to asses if symbionts of protists should be considered beneficial, neutral or detrimental for the host (Gast et al. 2009; Bernhard et al. 2018). Moreover, in some cases, the effect of the associated organism(s), whenever determined, has been observed to shift among the two poles, mutualism and parasitism, depending on several factors (e.g. host health, environmental changes, etc.) (Kusch et al. 2002; Hori and Fujishima 2003). Thus, considering all this theoretical and practical aspects, it seems appropriate to include also parasitism while defining the holobiont boundaries for taxonomic description of protists.

Last but not least, in case of protists the characterization of a host and its holobiome can be considered, in general, rather feasible in practice. Indeed, microorganisms associated with protists are usually few and, consequently, not so dramatically difficult to characterize, or at least, to detect. Moreover, obtaining protists’ mitogenomes is relatively straightforward, given the relative low complexity that is generally associated to the nuclear genome of protists (Lynch 2005; Gregory 2005; Haygood 2006), thus allowing to limit sequencing and analysis efforts.

The case study here proposed is fully representative. In the newly described ciliate species, *E. vanleeuwenhoeki* sp. nov., a sole, stably associated bacterium was detected, namely the endosymbiont “*Ca*. Pinguicoccus supinus”, which makes the application of the holobiont concept to species description unambiguous. As detailed above, the peculiar features of this symbiont, especially the genomic ones, are strongly indicative of a long coevolutionary and adaptation history with its host, so that this bacterium could be even hypothesized as an autoapomorphic character for the host species. Thus, this symbiotic system unambiguously fits the holobiont definition herein proposed for taxonomic purposes (highly integrated system with emerging properties).

In conclusion, in our view, the “next generation taxonomy” approach can constitute a new and valuable standard for unicellular eukaryotes descriptions and redescriptions. Indeed, integrating standard taxonomical analyses with the most modern tools in genomics and bioinformatics and the concept of holobiont, it provides additional descriptors valuable not only to fully and unambiguously define a species but also to infer its interaction with other organisms and, at least partially, its biology. At practical level, the suitability of the proposed approach for unicellular eukaryotic organisms, in particular free-living forms, associated with bacterial endobionts, is herein documented with the description of *E. vanleeuwenhoeki*, and we are rather confident that it should be similarly straightforward if applied to other kinds of holobiontic protists.

### Applicability of the “next generation taxonomy” to “higher-level” organisms

The hardest challenge for the “next generation taxonomy” is, for sure, to become suitable for “higher-level” organisms’ description/redescription. In this task there are, indeed, several issues. First of all, in general, there are some difficulties in defining which organisms may constitute a holobiont among multicellular and compartimentalized organisms such as metazoa (e.g. humans and associated gut microbiota), and therefore to determine if an associated organism should be included as an additional descriptor. Moreover, in this regard, the characteristics of the microbiota are also rather different in different types of organisms. In vertebrates, including humans, variations in the composition of the “holobiont”, even just focussing on the gut microbiota, are enormous. Microbiota composition varies during post-birth development, is profoundly influenced by the diet, and by a variety of environmental and genetic factors (Huse et al. 2012). Even a precise description of the core gut microbiota is problematic, and the focus is more toward the description of the core microbiome (Lloyd-Price et al. 2016). The scenario is similar in many arthropods, in particular in insects, where the composition of the microbial communities associated with the different body districts is highly variable (see for example Montagna et al. 2015; Montagna et al. 2016), with core microbiotas easily defined only in some taxa (e.g. in termites), or for some types of symbiotic associations (e.g. in the presence of vertically transmitted primary symbionts). Thus, while in dealing with well defined obligate/stable symbionts of insects (e.g. *Buchnera aphidicola*, “*Ca*. Zinderia insecticola”, etc.), the application of the holobiont concept to taxon description could be straightforward, it does not come easy when focussing on highly variable microbial communities. In any case, from a purely practical perspective, it would be not be feasible or even at all possible for researchers to identify and list all the possible associated microorganisms of the metazoans’ microbiota. Moreover, also the host’s mitogenome analysis could be not so straightforward, compared to unicellular organisms, given the size and the higher complexity of the nuclear genome of most metazoans (Lynch 2005; Gregory 2005; Haygood 2006). Even with the most modern sequencing and bioinformatic techniques available, this task could become too challenging and time-demanding for taxonomists.

In other words, from both a conceptual (i.e. how to apply the holobiont definition) and a practical (i.e. mitogenome and microbiome analyses) point of view, in our opinion, the present state of the art and technology do not allow to apply the “next generation taxonomy” approach to multicellular organisms.

To sum up, we consider the proposed workflow, which we named “next generation taxonomy”, extremely valuable and practicable for unicellular eukaryotic organisms, and we encourage the inclusion in their descriptions of mitogenome and associated bionts. On the other side, for multicellular organisms we consider this approach to be still premature, although attempts on more anatomically simple organisms (e.g. Placozoa (Driscoll et al. 2013), Orthonectida (Bondarenko et al. 2019)), with presumably lower complexity microbiomes, could be tempting.

## Nomenclature acts

The present work has been registered in ZooBank (code: LSID:urn:lsid:zoobank.org:pub:231DF703-DBA8-4839-9755-5C71D6872406), as well as the new *Euplotes* species, *Euplotes vanleeuwenhoeki* (code: LSID:urn:lsid:zoobank.org:act:19C2249B-BF2D-4E81-9391-A271D38CD7B6). The correspondent web pages are available at the following addresses: http://www.zoobank.org/References/231DF703-DBA8-4839-9755-5C71D6872406 and http://www.zoobank.org/NomenclaturalActs/19C2249B-BF2D-4E81-9391-A271D38CD7B6, respectively.

## Description of “*Candidatus* Pinguicoccus” gen. nov

*Pinguicoccus* [Pin.gui.coc’cus. L. adj. *pinguis* fat; N.L. n. *coccus* (from Gr. masc. n. *kokkos* grain, seed) coccus: N.L. masc. n. *Pinguicoccus*, a fat coccus, because its rounded body shape and because it was often found associated with host’s lipid droplets].

The genus description at present is the same as the description of the type species “*Ca.* Pinguicoccus supinus”.

## Description of “*Candidatus* Pinguicoccus supinus” sp. nov

*Pinguicoccus supinus* (su.pi’nus. L. adj. *supinus* lying on the back, supine, indolent, lazy, because it is not a motile microorganism and because it lacks several fundamental metabolic pathways).

Type species of the genus. Non motile *Verrucomicrobia* bacterium of family *Puniceicoccaceae* (class *Opitutae*). Roundish-ovoid in shape, detected in the cytoplasm of the ciliate *Euplotes vanleeuwenhoeki*, with a diameter of 1.3-2.3 µm. It usually lies beneath the ciliate cortex, often in clusters of several individuals. Cells are delimited by a double membrane with a thin space between the two layers; no symbiosome was ever observed. In several individuals, an invagination of the inner membrane and folding of the outer membrane are observed. Homogeneous bacterial cytoplasm. Sometimes shows presence of nucleoids. Reproduce by binary fission in the host cytoplasm, by means of an apparently typical symmetrical division. Its genome size is 163,218 bp and G+C content is 25%. The genetic code for protein translation is the “translation table 4”.

Base of attribution: 16S rRNA gene sequence, MK569697; complete genome sequence CP039370; recognized by oligonucleotidic probes EUB338 VII and Pingui_1174.

## Acknowledgements

The authors are grateful to the Marine Biology Laboratory, Andhra University, India, for providing the research facilities. In particular, Professor Kalavati, Professor Raman Akkur, and Professor Prabhakara Rao Yallapragada are acknowledged for their valuable aid. Special acknowledgements to Simone Gabrielli for tree and photo-editing support. This work was supported by the European Commission FP7-PEOPLE-2009-IRSES project CINAR PATHOBACTER (247658) (to GP); the University of Pisa PRA_2018_63 project (to GP); the Italian Ministry of Education, University and Research (MIUR): Dipartimenti di Eccellenza Program (2018–2022) - Dept. of Biology and Biotechnology “L. Spallanzani”, University of Pavia (to DS).

## Author Contributions

**Conceptualization:** GP, VS, LG
**Sampling and culturing:** VN, BVS
**Morphological and ultrastructural analyses:** VN, LM
**Molecular analyses:** VS, LG, VN
**Bioinformatics:** LG, MC, DS
**Phylogeny and phylogenomic analyses:** VS, LG, MC, GP
**Environmental screening on IMNGS:** OL
**Visualization:** VN, VS, LG, LM, OL, FV, GP
**Funding acquisition:** GP, FV, DS, CB
**Work supervision:** GP, LM
**Writing – original draft:** VS, LG, LM, OL
**Writing – review & editing:** GP, MC, DS, CB, VN, FV, BVS

