## Supplementary files 1-9 for "Next generation taxonomy: integrating traditional species description with the holobiont concept and genomic approaches - The in-depth characterization of a novel *Euplotes* species as a case study"

### SUPPLEMENTARY TABLES

**Supplementary Table 1.** List of primers used for *Euplotes vanleeuwenhoeki* sp. nov.18S rRNA gene sequencing

| Name | Sequence (5'-3') | Use | Type | Reference |
|---|---|---|---|---|
| 18S F9 | CTG GTT GAT CCT GCC AG | PCR | Forward | Medlin et al. 1988 |
| 18S R1513 Hypo | TGA TCC TTC YGC AGG TTC | PCR | Reverse | Petroni et al. 2002 |
| 18S R536 | CTG GAA TTA CCG CGG CTG | SEQ | Reverse | Rosati et al. 2004 |
| 18S R1052 | AAC TAA GAA CGG CCA TGC A | SEQ | Reverse | Rosati et al. 2004 |
| 18S F783 | GAC GAT CAG ATA CCG TC | SEQ | Forward | Rosati et al. 2004 |

*PCR* polymerase chain reaction*; SEQ* sequencing

**Supplementary Table 2.** List of highly reduced bacterial genomes used for COGs analysis

| Species | Accession Number | Taxonimic | Length (bp) | Number of proteins | Numer of COGs |
|---|---|---|---|---|---|
| *“Candidatus* Carsonella ruddii*”* | CP024798 | *γ-Proteobacteria* | 174,004 | 200 | 157 |
| *“Candidatus* Hodgkinia cicadicola*”* | CP001226 | *α-Proteobacteria* | 143,795 | 170 | 132 |
| *“Candidatus* Sulcia muelleri*”* | CP016223 | *Bacteroidetes* | 192,244 | 190 | 179 |
| *“Candidatus* Tremblaya princeps*”* | LN999057 | *β-Proteobacteria* | 143,340 | 116 | 109 |
| *“Candidatus* Tremblaya phanecola*”* | CP003982 | *β-Proteobacteria* | 171,500 | 162 | 174 |
| *“Candidatus* Zinderia insecticola*”* | CP002161 | *β-Proteobacteria* | 208,564 | 206 | 185 |
| *“Candidatus* Nasuia deltocephalinicola*”* | CP013211 | *β-Proteobacteria* | 112,031 | 141 | 106 |

**Supplementary Table 3.** Selected complete genomes of *Verrucomicrobia* used for COGs analysis

| Species | Accession Number | Number of COGs | COGs shared with "*Ca.* Pinguicoccus supinus" | Uniques COGs |
|---|---|---|---|---|
| *Methylacidiphilum infernorum* strain V4 | GCA_000019665 | 1178 | 127 | 15 |
| *Methylacidiphilum fumariolicum* strain SolIV | GCA_000019665 | 1187 | 126 | 22 |
| *Akkermansia glycaniphila* | GCA_900097105 | 1235 | 123 | 41 |
| *Akkermansia muciniphila* strain ATCC | GCA_000020225 | 1192 | 124 | 5 |
| *Akkermansia muciniphila* strain YL44 | GCA_001688765 | 1233 | 123 | 12 |
| *Coraliomargarita akajimensis* | GCA_00025905 | 1463 | 128 | 68 |
| *Opitutus terrae* strain PB90-1 | GCA_000019965 | 1750 | 130 | 91 |
| *Lacunisphaera limnophila* | GCA_001746835 | 1564 | 127 | 48 |
| "*Candidatus* Xiphinematobacter" sp. Idaho Grape | GCA_001318295 | 652 | 123 | 10 |
| Uncultured bacterium strain IMCC | GCA_000972765 | 1399 | 125 | 30 |
| Uncultured bacterium strain HZ-65 | GCA_002310495 | 1630 | 35 | 48 |
| Uncultured bacterium strain TAV5 | GCA_002310495 | 1769 | 128 | 128 |
| "*Candidatus* Pinguicoccus supinus" | CP039370 | 133 | 133 | 0 |

**Supplementary Table 4.** Sequences belonging to the Superphylum PVC, not shown in the *Verrucomicrobia* phylogenetic tree (Figure 7)

| **Species** | **Accession Number** |
|---|---|
| *Cerasicoccus arenae* | AB292183 |
| *Cerasicoccus maritimus* | AB372849 |
| *Cerasicoccus frondis* | NR_112768 |
| *Pelagicoccus albus* | AB286016 |
| *Pelagicoccus litoralis* | AB286017 |
| *Pelagicoccus mobilis* | AB286015 |
| *Pelagicoccus croceus* | AB297922 |
| *Rubritalea profundi* | KR108285 |
| *Rubritalea tangerina* | AB297806 |
| *Rubritalea sabuli* | AB353310 |
| *Rubritalea squalenifaciens* | AB277853 |
| *Roseibacillus ishigakijimensis* | AB331888 |
| *Roseibacillus ponti* | AB331889 |
| *Roseibacillus persicicus* | AB331892 |
| *Haloferula harenae* | AB372852 |
| *Haloferula rosea* | AB372853 |
| *Haloferula helveola* | AB372855 |
| *Haloferula sargassicola* | AB372856 |
| *Prosthecobacter algae* | NR_133826 |
| *Prosthecobacter debontii* | AJ966882 |
| *Prosthecobacter fluviatilis* | AB305640 |
| *Prosthecobacter vanneervenii* | AJ966883 |
| *Brevifollis gellanilyticus* | AB552872 |
| *Methylacidimicrobium cyclopophantes* | KM210555 |
| *Methylacidimicrobium fagopyrum* | KM210553 |
| *Methylacidimicrobium tartarophylax* | KM210554 |
| *Lentisphaera araneosa* | ABCK01000003 |
| *Lentisphaera marina* | JN175275 |

**Supplementary Table 5.** Identity values among 18S rRNA sequences of selected *Euplotes,* phylogenetically close to *Euplotes vanleeuwenhoeki* sp. nov.

| | a. | b. | c. | d. | e. | **f.** | g. | h. | i. | j. | k. | l. | m. | n. | o. | p. | q. |
|---|---|---|---|---|---|---|---|---|---|---|---|---|---|---|---|---|---|
| a. *Euplotes charon,* AF492705 | - | | | | | | | | | | | | | | | | |
| b. *E. magnicirratus,* AJ549210 | 100 | - | | | | | | | | | | | | | | | |
| c. *E. euryhalinus,* EF094968 | 97.8 | 97.8 | - | | | | | | | | | | | | | | |
| d. *E. euryhalinus,* JF903799 | 96.9 | 96.9 | 98.6 | - | | | | | | | | | | | | | |
| e. *E. trisulcatus,* EF690810 | 95.7 | 95.8 | 96.9 | 96.2 | - | | | | | | | | | | | | |
| **f. *E. vanleeuwenhoeki,* KY855568** | **96.1** | **96.2** | **97.2** | **96.6** | **98.7** | - | | | | | | | | | | | |
| g. *E.* cf. *antarcticus,* FJ998023 | 96.1 | 96.2 | 96.9 | 96.6 | 98.6 | **99.0** | - | | | | | | | | | | |
| h. *E. rariseta,* JX437135 | 96.7 | 96.8 | 97.4 | 96.9 | 95.6 | **96.3** | 96.2 | - | | | | | | | | | |
| i. *E. rariseta,* AJ305248 | 96.7 | 96.7 | 97.3 | 96.9 | 95.6 | **96.3** | 96.2 | 100 | - | | | | | | | | |
| j. *E. charon,* JF694043 | 95.8 | 95.9 | 96.6 | 96.1 | 94.9 | **95.4** | 95.4 | 96.2 | 96.2 | - | | | | | | | |
| k. *E. parkei,* AJ305247 | 96.1 | 96.1 | 96.7 | 96.3 | 95.0 | **95.6** | 95.6 | 96.5 | 96.4 | 99.6 | - | | | | | | |
| l. *E. focardii,* EF094961 | 96.1 | 96.2 | 96.4 | 95.9 | 94.7 | **95.5** | 95.4 | 96.2 | 96.2 | 98.5 | 98.8 | - | | | | | |
| m. *E. quinquecarinatus,* JX437136 | 96.2 | 96.3 | 96.5 | 96.1 | 94.7 | **95.4** | 95.3 | 96.5 | 96.5 | 98.9 | 99.3 | 99.0 | - | | | | |
| n. *E. vannus,* AJ305241 | 95.9 | 95.9 | 96.1 | 95.8 | 95.2 | **95.7** | 96.0 | 96.0 | 96.0 | 96.9 | 97.1 | 97.0 | 97.1 | - | | | |
| o. *E. crassus,* AJ305239 | 95.8 | 95.9 | 96.2 | 95.7 | 95.3 | **95.9** | 95.9 | 96.0 | 95.9 | 96.7 | 96.8 | 96.8 | 96.8 | 99.0 | - | | |
| p. *E. cristatus,* GU953667 | 96.1 | 96.2 | 96.3 | 95.9 | 95.6 | **96.1** | 96.3 | 96.4 | 96.4 | 97.1 | 97.4 | 97.1 | 97.4 | 99.2 | 98.9 | - | |
| q. *E. minuta,* AJ305244 | 96.3 | 96.4 | 96.4 | 96.0 | 95.6 | **96.2** | 96.3 | 96.5 | 96.5 | 97.4 | 97.5 | 97.1 | 97.1 | 98.3 | 97.9 | 98.8 | - |

Sequence obtained in the present work is shown in *bold.*

**Supplementary Table 6.** List of contigs in the preliminary assembly firstly annotated as bacterial

| Node | Node lenght (bp) | Coverage | Predicted ORF name | Best blast hit | Best hit evalue | Best hit species | Best hit kingdom |
|---|---|---|---|---|---|---|---|
| NODE_63513_length_223_cov_33.8014 | 223 | 55.40 | PROKKA_00150 | gb\|AHM82281.1\| Antirestriction protein klcA (plasmid) | 7E-14 | *Klebisella pneumonie* | Bac |
| NODE_54228_length_256_cov_9.56983 | 256 | 23.62 | PROKKA_00100 | emb\|CUJ48696.1\| Uncharacterised protein | 0,000009 | *Achromobacter sp.* | Bac |
| NODE_54133_length_257_cov_25.6667 | 257 | 41.95 | | | | | NA |
| NODE_39611_length_344_cov_7701.77 | 344 | 545.73 | | | | | Bac |
| NODE_33671_length_401_cov_8124.2 | 401 | 616.19 | | | | | NA |
| NODE_26446_length_512_cov_14.6124 | 512 | 41.88 | | | | | NA |
| NODE_23590_length_581_cov_19691.1 | 581 | 4038.93 | PROKKA_00050 | emb\|CQD05533.1\| Uncharacterised protein | 0,000000001 | *Wolbachia endosymbiont* | Bac |
| NODE_20813_length_663_cov_12.8225 | 663 | 33.01 | | | | | NA |
| NODE_20354_length_681_cov_5597.38 | 681 | 2656.65 | PROKKA_00047 | gb\|ACX30947.1\| cytochrome c oxidase subunit 2 | 0,0005 | *Monoeuplotes minuta* | Euk |
| NODE_17358_length_812_cov_16.698 | 812 | 42.15 | | | | | NA |
| NODE_14225_length_995_cov_16.4281 | 995 | 45.03 | | | | | NA |
| NODE_13189_length_1068_cov_102.537 | 1068 | 210.66 | PROKKA_00009 | gb\|KPY54789.1\| Peroxisomal catalase | 6E-27 | *Pseudomonas syringae* | Bac |
| | | | PROKKA_00010 | gb\|OMJ82787.1\| hypothetical protein SteCoe_16437 | 2E-11 | *Stentor coeruleus* | Euk |
| NODE_11646_length_1204_cov_18.6664 | 1204 | 47.12 | PROKKA_00008 | ref\|WP_053224099.1\| hypothetical protein | 3E-75 | *Roseivirga seohaensis* | Bac |
| NODE_9993_length_1241_cov_21.5425 | 1241 | 54.81 | PROKKA_00266 | emb\|CDW87446.1\| 2-oxoglutarate dehydrogenase | 1E-38 | *Stylonychia lemnae* | Euk |
| NODE_10734_length_1306_cov_14.3987 | 1306 | 38.71 | PROKKA_00007 | ref\|WP_073123159.1\| hypothetical protein | 4E-13 | *Reichenbachiella agariperforans* | Bac |
| NODE_10516_length_1331_cov_8909.78 | 1331 | 1026.9 | | | | | NA |
| NODE_10124_length_1378_cov_26.7218 | 1378 | 63.07 | PROKKA_00001 | dbj\|BAF01922.1\| 2-oxoglutarate dehydrogenase, E1 compone | 1E-145 | *Arabidopsis thaliana* | Euk |
| NODE_9920_length_1405_cov_8414.95 | 1405 | 857.98 | | | | | NA |
| NODE_9773_length_1425_cov_8373.22 | 1425 | 1105.09 | | | | | NA |
| NODE_8563_length_1620_cov_15.2618 | 1620 | 40.16 | PROKKA_00257 | gb\|EJY82326.1\| Translation initiation factor IF-2 | 7E-61 | *Oxytricha trifallax* | Euk |
| NODE_7688_length_1800_cov_7630.52 | 1800 | 1104.66 | | | | | NA |
| NODE_7375_length_1868_cov_7420.38 | 1868 | 1089.09 | | | | | NA |
| NODE_6901_length_1971_cov_17.3391 | 1971 | 44.82 | | | | | NA |

| | | | | | | | |
|---|---|---|---|---|---|---|---|
| NODE_6789_length_2020_cov_7470.16 | 2020 | 1218.48 | | | | | NA |
| NODE_6787_length_2021_cov_7876.26 | 2021 | 1515.81 | | | | | NA |
| NODE_6776_length_2023_cov_7459.12 | 2023 | 1163.49 | | | | | NA |
| NODE_6775_length_2024_cov_7520.83 | 2024 | 1191.06 | | | | | NA |
| NODE_6647_length_2066_cov_7559.65 | 2066 | 1339.86 | | | | | NA |
| NODE_6597_length_2080_cov_7428.11 | 2080 | 1969.48 | | | | | NA |
| NODE_6377_length_2152_cov_8030.21 | 2152 | 1259.42 | | | | | NA |
| NODE_6321_length_2170_cov_7456.77 | 2170 | 2459.42 | | | | | NA |
| NODE_6187_length_2210_cov_22.5391 | 2210 | 50.65 | PROKKA_00141 | emb\|CDW75606.1\| dihydrolipoyl dehydrogenase | 0 | *Stylonychia lemnae* | Euk |
| NODE_6090_length_2245_cov_41.5203 | 2245 | 69.48 | PROKKA_00139 | gb\|OGN76630.1\| bifunctional methylenetetrahydrofolate | 1E-50 | *Cloroflexi bacterium* | Bac |
| NODE_4338_length_3087_cov_29.2668 | 3087 | 65.76 | PROKKA_00081 | emb\|CDW90485.1\| s-adenosylmethionine synthetase | 7E-71 | *Stylonychia lemnae* | Euk |
| | | | PROKKA_00082 | gb\|EJY72651.1\| S-adenosylmethionine synthase | 1E-65 | *Oxytricha trifallax* | Euk |
| | | | PROKKA_00083 | emb\|CDW81045.1\| probable splicing factor 3a subunit 1-li | 4E-28 | *Stylonychia lemnae* | Euk |
| NODE_3383_length_3801_cov_31.3008 | 3801 | 69.26 | PROKKA_00063 | gb\|AOQ25829.1\| ABC transporter B family member 9-like pr | 0 | *Euplotes crassus* | Euk |
| NODE_3264_length_3917_cov_35.7279 | 3917 | 81.87 | | | | | NA |
| NODE_2918_length_4316_cov_24.9868 | 4316 | 58.42 | PROKKA_00053 | gb\|EJY82189.1\| Citrate synthase | 8E-125 | *Oxytricha Trifallax* | Euk |
| | | | PROKKA_00054 | emb\|CDW91737.1\| ubiquitin-conjugating enzyme family prot | 1E-33 | *Stylonychia lemnae* | Euk |
| | | | PROKKA_00056 | emb\|CDW91737.1\| ubiquitin-conjugating enzyme family prot | 0 | *Stylonychia lemnae* | Euk |
| NODE_1691_length_6468_cov_35.0074 | 6468 | 58.78 | PROKKA_00036 | emb\|CDW89925.1\| aminopeptidase n | 4E-65 | *Stylonychia lemnae* | Euk |
| | | | PROKKA_00037 | ref\|WP_052469931.1\| aminopeptidase N | 3E-25 | *Thiolapillus brandeum* | Bac |
| | | | PROKKA_00041 | ref\|WP_035961980.1\| aminopeptidase N | 1E-14 | *Kocuria marina* | Bac |
| | | | PROKKA_00042 | ref\|WP_063796675.1\| aminopeptidase N | 0,0000001 | *Chondromyces crocatus* | Bac |
| | | | PROKKA_00043 | ref\|WP_006288812.1\| aminopeptidase N | 9E-17 | *Parascardovia denticolens* | Bac |
| NODE_1576_length_6757_cov_43.8379 | 6757 | 100.74 | PROKKA_00026 | ref\|XP_003061505.1\| predicted protein | 1E-74 | *Micromonas pusilla* | Bac |
| | | | PROKKA_00027 | gb\|EJY81362.1\| hypothetical protein OXYTRI_21126 | 5E-39 | *Oxytricha Trifallax* | Euk |
| | | | PROKKA_00028 | emb\|CDW78435.1\| asparaginyl-trna synthetase | 8E-38 | *Stylonychia lemnae* | Euk |

| | | | | | | | |
|---|---|---|---|---|---|---|---|
| | | | PROKKA_00029 | ref\|WP_025644854.1\| MULTISPECIES: valine--tRNA ligase | 0,00005 | *Psychrobacter* | Bac |
| | | | PROKKA_00031 | ref\|XP_016588471.1\| valyl-tRNA synthetase | 1E-12 | *Sporothrix schenckii* | Euk |
| | | | PROKKA_00032 | gb\|OMJ78400.1\| hypothetical protein SteCoe_21773 | 2E-59 | *Stentor coeruleus* | Euk |
| | | | PROKKA_00033 | ref\|XP_002963439.1\| hypothetical protein SELMODRAFT_1419 | 9E-35 | *Selaginella moellendorffii* | Euk |
| | | | PROKKA_00034 | ref\|XP_013872938.1\| PREDICTED: acylamino-acid-releasing | 6E-10 | *Austrofundulus limnaeus* | Euk |
| NODE_1359_length_7366_cov_33.3603 | 7366 | 82.13 | | | | | NA |
| NODE_793_length_9898_cov_33.6908 | 9898 | 76.94 | PROKKA_00247 | gb\|EJY85937.1\| GTPase | 8E-106 | *Oxytricha trifallax* | Euk |
| | | | PROKKA_00249 | emb\|CDW88205.1\| UNKNOWN | 0,00002 | *Stylonychia lemnae* | Euk |
| | | | PROKKA_00254 | ref\|XP_001447435.1\| hypothetical protein | 1E-13 | *Paramecium tetraurelia* | Euk |
| | | | PROKKA_00255 | gb\|KRX04237.1\| hypothetical protein PPERSA_11361 | 1E-82 | *Pseudocohnilembus persalinus* | Euk |
| NODE_713_length_10472_cov_30.1492 | 10472 | 66.98 | PROKKA_00219 | gb\|EJY65049.1\| Putative non-transporter ABC protein | 6E-129 | *Oxytricha trifallax* | Euk |
| | | | PROKKA_00224 | emb\|CDW89759.1\| adenylosuccinate lyase | 2E-167 | *Stylonychia lemnae* | Euk |
| | | | PROKKA_00226 | gb\|EJY78390.1\| hypothetical protein OXYTRI_24455 | 1E-16 | *Oxytricha trifallax* | Euk |
| | | | PROKKA_00227 | emb\|CDW91341.1\| UNKNOWN | 4E-33 | *Stylonychia lemnae* | Euk |
| | | | PROKKA_00231 | emb\|CDW86186.1\| histidine acid phosphatase family protein | 9E-11 | *Stylonychia lemnae* | Euk |
| NODE_652_length_10908_cov_32.696 | 10908 | 74.56 | PROKKA_00166 | gb\|EJY87052.1\| hypothetical protein OXYTRI_07502 | 1E-142 | *Oxytricha trifallax* | Euk |
| NODE_567_length_11743_cov_63.0045 | 11743 | 141.53 | PROKKA_00119 | ref\|XP_003385623.1\| PREDICTED: 4-aminobutyrate aminotran | 3E-50 | *Amphimedon queenslandica* | Euk |
| | | | PROKKA_00120 | ref\|XP_012234736.1\| PREDICTED: aspartate--tRNA ligase, c | 8E-62 | *Linepithema humile* | Euk |
| | | | PROKKA_00121 | gb\|EJY72918.1\| Aspartyl-tRNA synthetase | 1E-85 | *Oxytricha trifallax* | Euk |
| | | | PROKKA_00122 | gb\|ODA77684.1\| hypothetical protein RJ55_06286 | 3E-55 | *Drechmeria coniospora* | Euk |
| | | | PROKKA_00123 | emb\|CDW78194.1\| UNKNOWN | 0,000000006 | *Stylonychia lemnae* | Euk |
| | | | PROKKA_00136 | gb\|OMJ81750.1\| hypothetical protein SteCoe_17746 | 0,00001 | *Stentor coeruleus* | Euk |

| | | | | | | | |
|---|---|---|---|---|---|---|---|
| NODE_549_length_11894_cov_125.266 | 11894 | 28.07 | PROKKA_00101 | emb\|CDW74791.1\| UNKNOWN | 3E-22 | *Stylonychia lemnae* | Euk |
| | | | PROKKA_00107 | gb\|EJY69552.1\| hypothetical protein OXYTRI_09710 | 3E-23 | *Oxytricha trifallax* | Euk |
| | | | PROKKA_00109 | gb\|EJY74005.1\| hypothetical protein OXYTRI_04742 | 0,00000007 | *Oxytricha trifallax* | Euk |
| | | | PROKKA_00111 | gb\|EJY69380.1\| hypothetical protein OXYTRI_10000 | 2E-41 | *Oxytricha trifallax* | Euk |
| | | | PROKKA_00112 | gb\|EJY81593.1\| hypothetical protein OXYTRI_20893 | 3E-41 | *Oxytricha trifallax* | Euk |
| | | | PROKKA_00113 | gb\|EJY65673.1\| hypothetical protein OXYTRI_14171 | 8E-45 | *Oxytricha trifallax* | Euk |
| | | | PROKKA_00114 | gb\|EJY78913.1\| hypothetical protein OXYTRI_23921 | 0,0000003 | *Oxytricha trifallax* | Euk |
| | | | PROKKA_00117 | gb\|EJY82990.1\| Ribonucleoside-diphosphate reductase | 0 | *Oxytricha trifallax* | Euk |
| | | | PROKKA_00118 | ref\|XP_012652545.1\| ribonucleoside-diphosphate reductase | 3E-79 | *Tetrahymena termophila* | Euk |
| NODE_522_length_12235_cov_34.7649 | 12235 | 80.91 | PROKKA_00086 | gb\|EJY71120.1\| Fumarate hydratase | 0 | *Oxytricha Trifallax* | Euk |
| | | | PROKKA_00087 | ref\|XP_001022217.2\| rhodanese-like domain protein | 4E-26 | *Tetrahymena termophila* | Euk |
| | | | PROKKA_00088 | gb\|EJY86417.1\| hypothetical protein OXYTRI_15059 | 0,0001 | *Oxytricha Trifallax* | Euk |
| | | | PROKKA_00093 | emb\|CDW88321.1\| UNKNOWN | 0,000004 | *Stylonychia lemnae* | Euk |
| | | | PROKKA_00095 | emb\|CDW88321.1\| UNKNOWN | 6E-14 | *Stylonychia lemnae* | Euk |
| | | | PROKKA_00096 | emb\|CDW88321.1\| UNKNOWN | E-31 | *Stylonychia lemnae* | Euk |
| NODE_426_length_13579_cov_33.0478 | 13579 | 78.62 | PROKKA_00071 | gb\|AKJ66198.1\| casein kinase II beta-1 (macronuclear) | 2E-70 | *Euplotes octarinatus* | Euk |
| | | | PROKKA_00074 | emb\|CDW76091.1\| short-chain dehydrogenase | 2E-23 | *Chryseolinea serpens* | Bac |
| | | | PROKKA_00080 | emb\|CEL97133.1\| unnamed protein product | 1E-70 | *Vitrella brassicaformis* | Euk |
| | | | PROKKA_00060 | emb\|CDW89135.1\| adenylosuccinate synthetase | 9E-112 | *Stylonychia lemnae* | Euk |
| | | | PROKKA_00168 | ref\|WP_073141524.1\| glycine dehydrogenase | 2E-130 | *Chryseolinea serpens* | Bac |
| | | | PROKKA_00169 | gb\|EJY88375.1\| hypothetical protein OXYTRI_16562 | 0 | *Oxytricha Trifallax* | Euk |

*Red* colour: bacterial ORF; *Green:* eukaryotic ORF; *Blue:* not annotated ORF.

**Supplementary Table 7.** Identity values among 16S rRNA sequences of selected *Opitutae* bacteria, phylogenetically close to "*Candidatus* Pinguicoccus supinus"

| | a. | b. | c. | d. | e. | f. | g. | h. | i. | j. | k. | l. | m. | **n.** | o. | p. | q. | r. | s. | t. | u. | v. | w. | x. | y. |
|---|---|---|---|---|---|---|---|---|---|---|---|---|---|---|---|---|---|---|---|---|---|---|---|---|---|
| a. *Coraliomargarita akajimensis,* CP001998 | - | | | | | | | | | | | | | | | | | | | | | | | | |
| b. *Fucophilus fucoidanolyticus,* AB073978 | 94.4 | - | | | | | | | | | | | | | | | | | | | | | | | |
| c. *Puniceicoccus vermicola,* DQ539046 | 88.5 | 87.9 | - | | | | | | | | | | | | | | | | | | | | | | |
| d. *Cerasicoccus frondis,* NR_112768 | 87.4 | 87.4 | 88.0 | - | | | | | | | | | | | | | | | | | | | | | |
| e. *Ruficoccus amylovorans,* KT751307 | 88.3 | 86.9 | 87.3 | 90.0 | - | | | | | | | | | | | | | | | | | | | | |
| f. Epixenosome, Y19169 | 87.1 | 87.6 | 85.6 | 86.4 | 85.9 | - | | | | | | | | | | | | | | | | | | | |
| g. Unc. rumen bact., AB614893 | 82.7 | 82.5 | 81.2 | 81.1 | 82.4 | 80.0 | - | | | | | | | | | | | | | | | | | | |
| h. Unc. rumen bact., AB034150 | 82.4 | 82.5 | 81.0 | 81.0 | 82.0 | 80.2 | 97.8 | - | | | | | | | | | | | | | | | | | |
| i. Unc. rumen bact., AB615161 | 82.5 | 82.2 | 80.0 | 80.5 | 81.7 | 79.3 | 90.3 | 90.4 | - | | | | | | | | | | | | | | | | |
| j. Unc. bact., HQ155682 | 84.3 | 83.7 | 82.0 | 82.6 | 84.0 | 81.1 | 87.5 | 86.9 | 86.7 | - | | | | | | | | | | | | | | | |
| k. Unc. rumen bact., EU850497 | 84.3 | 83.3 | 81.5 | 81.9 | 83.6 | 79.5 | 87.8 | 87.7 | 87.5 | 94.7 | - | | | | | | | | | | | | | | |
| l. Unc. bact., AY571501 | 85.0 | 85.0 | 82.1 | 83.0 | 84.1 | 82.4 | 85.8 | 85.7 | 86.4 | 88.9 | 90.2 | - | | | | | | | | | | | | | |
| m. Unc. bact., JQ993517 | 85.1 | 85.1 | 82.0 | 82.9 | 84.1 | 82.3 | 85.9 | 85.8 | 86.4 | 89.0 | 90.3 | 100 | - | | | | | | | | | | | | |
| **n. "*Ca.* Pinguicoccus supinus"** | **75.9** | **76.0** | **75.6** | **75.6** | **76.8** | **76.4** | **75.7** | **76.1** | **76.3** | **77.1** | **76.9** | **78.2** | **78.3** | - | | | | | | | | | | | |
| o. Verrucomicrobia bact., MNWT01000005 | 87.4 | 86.4 | 87.1 | 87.9 | 88.8 | 86.4 | 84.2 | 84.3 | 84.4 | 84.5 | 85.5 | 86.4 | 86.4 | **78.8** | - | | | | | | | | | | |
| p. Unc. bact., JQ993599 | 85.8 | 85.8 | 84.1 | 85.3 | 85.8 | 84.5 | 82.9 | 82.9 | 82.0 | 82.7 | 82.5 | 84.7 | 84.8 | **78.1** | 87.6 | - | | | | | | | | | |
| q. Unc. bact., JQ993626 | 86.0 | 86.1 | 84.4 | 85.5 | 86.0 | 84.7 | 82.9 | 83.2 | 82.1 | 82.8 | 82.6 | 84.8 | 84.9 | **78.1** | 87.9 | 99.8 | - | | | | | | | | |
| r. Unc. bact., JQ993596 | 85.8 | 85.9 | 84.2 | 85.3 | 85.9 | 84.6 | 83.0 | 83.2 | 82.0 | 82.7 | 82.6 | 84.6 | 84.7 | **77.9** | 87.9 | 99.5 | 99.7 | - | | | | | | | |
| s. Unc. bact., AB826704 | 85.8 | 85.7 | 84.4 | 85.3 | 85.8 | 84.4 | 82.3 | 82.7 | 81.7 | 82.6 | 82.2 | 84.1 | 84.2 | **78.0** | 86.9 | 97.4 | 97.7 | 97.4 | - | | | | | | |
| t. Unc. bact., AB198611 | 85.7 | 86.9 | 83.7 | 85.3 | 86.7 | 84.3 | 82.8 | 82.7 | 81.9 | 83.3 | 83.3 | 85.3 | 85.4 | **77.7** | 87.1 | 90.8 | 91.0 | 91.2 | 90.3 | - | | | | | |
| u. Unc. bact., AB826705 | 85.8 | 86.0 | 85.2 | 85.9 | 86.4 | 84.2 | 83.5 | 83.5 | 82.8 | 83.2 | 83.2 | 84.8 | 84.9 | **78.3** | 87.2 | 91.4 | 91.7 | 91.8 | 90.8 | 95.0 | - | | | | |
| v. Unc. bact., GU472738 | 83.3 | 84.3 | 81.1 | 83.9 | 84.5 | 81.7 | 80.4 | 80.3 | 81.0 | 82.9 | 82.8 | 83.1 | 83.1 | **73.5** | 85.2 | 83.3 | 83.5 | 83.4 | 83.0 | 83.2 | 82.6 | - | | | |
| w. Unc. bact., AB231044 | 83.7 | 83.4 | 81.5 | 82.9 | 84.6 | 81.8 | 80.5 | 80.8 | 80.7 | 82.1 | 81.8 | 82.4 | 82.4 | **74.5** | 83.1 | 83.0 | 83.2 | 83.2 | 82.7 | 83.1 | 82.8 | 88.5 | - | | |
| x. Unc. bact., EU462461 | 83.7 | 83.8 | 82.1 | 83.3 | 84.2 | 82.4 | 80.5 | 80.5 | 81.0 | 80.7 | 80.2 | 81.7 | 81.7 | **74.8** | 83.3 | 83.5 | 83.7 | 83.7 | 83.4 | 82.6 | 83.0 | 85.7 | 86,1 | - | |
| y. *Pelagicoccus albus,* AB286016 | 86.2 | 85.9 | 85.3 | 86 | 86.7 | 85.7 | 80.9 | 80.8 | 80.7 | 81.8 | 81.3 | 82.6 | 82.6 | **75.4** | 86.6 | 84.9 | 85.1 | 84.9 | 84.9 | 85.3 | 86.3 | 82.4 | 82,5 | 82,5 | - |

Unc.: ucultured; bact.: bacterium. Sequence of "*Ca.* Pinguicoccus supinus" (accession number: MK569697) obtained in the present work is shown in *bold.* Square indicates sequences in the same clade of "*Ca.* Pinguicoccus supinus", in the phylogenetical analysis (see Figure 7).

**Supplementary Table 8.** Number of COGs in each retrieved category for the endosymbiont and the bacteria with highly reduced genome

| **Species** | **C** | **E** | **D** | **G** | **F** | **I** | **H** | **K** | **J** | **M** | **L** | **O** | **Q** | **P** | **S** | **R** | **T** | **U** | **V** |
|---|---|---|---|---|---|---|---|---|---|---|---|---|---|---|---|---|---|---|---|
| “*Candidatus* Carsonella ruddii” | 11 | 52 | 1 | 6 | 7 | 1 | 5 | 5 | 64 | 1 | 6 | 10 | 0 | 1 | 0 | 0 | 1 | 0 | 1 |
| “*Ca.* Hodgkinia cicadicola” | 11 | 18 | 0 | 1 | 2 | 0 | 18 | 5 | 63 | 0 | 2 | 10 | 1 | 2 | 1 | 2 | 0 | 0 | 0 |
| “*Ca.* Sulcia muelleri” | 17 | 47 | 1 | 3 | 3 | 1 | 7 | 6 | 80 | 3 | 5 | 15 | 1 | 1 | 1 | 1 | 0 | 4 | 1 |
| “*Ca.* Pinguibacter supinus” | 8 | 2 | 0 | 3 | 0 | 12 | 2 | 7 | 71 | 12 | 8 | 17 | 4 | 1 | 0 | 3 | 1 | 0 | 0 |
| “*Ca.*Zinderia insecticola” | 23 | 28 | 1 | 0 | 3 | 2 | 16 | 7 | 87 | 0 | 14 | 11 | 1 | 3 | 0 | 3 | 0 | 1 | 0 |
| “*Ca.*Tremblaya princeps” | 1 | 22 | 0 | 1 | 3 | 0 | 3 | 5 | 61 | 1 | 6 | 10 | 0 | 1 | 1 | 2 | 0 | 0 | 0 |
| “*Ca.* Tremblaya phenacola” | 4 | 44 | 0 | 2 | 6 | 0 | 5 | 5 | 85 | 2 | 5 | 11 | 0 | 1 | 0 | 2 | 1 | 0 | 1 |
| “*Ca.* Nasuia deltocephalinicola” | 7 | 14 | 0 | 1 | 1 | 0 | 6 | 6 | 54 | 1 | 4 | 12 | 0 | 1 | 0 | 1 | 0 | 0 | 0 |

**Supplementary Table 9.** List of endosymbiont positive samples retrieved during environmental screening in IMNGS

| #SampleID | Description | Total sequences | 16S rRNA gene similarity threshold | | | % Abundance |
|---|---|---|---|---|---|---|
| | | | 0.99 | 0.97 | 0.95 | |
| DRR016801 | Shrimp | 211797 | 3 | 3 | 3 | 0,001 |
| DRR092437 | Shrimp | 16878 | 1 | 1 | 1 | 0,006 |
| DRR092441 | Shrimp | 20252 | 1 | 1 | 1 | 0,005 |
| DRR092446 | Shrimp | 17090 | 1 | 1 | 1 | 0,006 |
| DRR092447 | Shrimp | 15391 | 1 | 1 | 1 | 0,006 |
| ERR1552052 | Microbial mat | 93189 | 0 | 1 | 1 | 0,001 |
| ERR1552095 | Microbial mat | 80377 | 0 | 3 | 3 | 0,004 |
| ERR1552096 | Microbial mat | 89254 | 0 | 1 | 1 | 0,001 |
| ERR1552097 | Microbial mat | 77545 | 0 | 1 | 1 | 0,001 |
| ERR1552105 | Microbial mat | 79413 | 0 | 0 | 1 | 0,001 |
| ERR1552200 | Microbial mat | 114958 | 0 | 0 | 2 | 0,002 |
| ERR1552219 | Microbial mat | 93938 | 0 | 1 | 1 | 0,001 |
| ERR1552221 | Microbial mat | 118382 | 0 | 0 | 1 | 0,001 |
| ERR1552229 | Microbial mat | 110619 | 0 | 1 | 1 | 0,001 |
| ERR1552230 | Microbial mat | 114188 | 0 | 3 | 3 | 0,003 |
| ERR1552231 | Microbial mat | 113875 | 0 | 1 | 1 | 0,001 |
| ERR1552232 | Microbial mat | 114328 | 0 | 1 | 1 | 0,001 |
| ERR1552233 | Microbial mat | 111589 | 0 | 1 | 1 | 0,001 |
| ERR1552241 | Microbial mat | 100058 | 0 | 0 | 1 | 0,001 |
| ERR1552251 | Microbial mat | 105726 | 0 | 0 | 1 | 0,001 |
| ERR1552265 | Microbial mat | 131893 | 0 | 1 | 1 | 0,001 |
| ERR1552276 | Microbial mat | 120044 | 0 | 1 | 1 | 0,001 |
| ERR1894936 | Seawater | 612186 | 50 | 50 | 50 | 0,008 |
| ERR1894938 | Seawater | 541615 | 2 | 2 | 2 | 0,000 |
| ERR574411 | Wastewater | 102263 | 0 | 5 | 5 | 0,005 |
| **ERR574415** | **Wastewater** | **125453** | **827** | **931** | **944** | **0,752** |
| ERR574416 | Wastewater | 99118 | 7 | 14 | 14 | 0,014 |
| **ERR574419** | **Wastewater** | **100345** | **83** | **115** | **116** | **0,116** |
| SRR2033822 | Plant | 9272 | 1 | 1 | 1 | 0,011 |
| SRR3169811 | Soil | 67644 | 1 | 1 | 1 | 0,001 |
| SRR3173821 | Soil | 57162 | 1 | 1 | 1 | 0,002 |
| SRR3173823 | Soil | 67517 | 1 | 1 | 1 | 0,001 |

For each positive sample are reported the SampleID, description, total number of sequences, positive hits with diverse similarity thresholds (99, 97, 95%), and the abundance percentage (i.e. the ratio between positive hits and total number of sequences). In bold are reported the percentages of the most abundant samples.
